# The roles of isolation and interspecific interaction in generating the functional diversity of an insular mammal radiation

**DOI:** 10.1101/2022.07.15.500274

**Authors:** Jonathan A. Nations, Brooks A. Kohli, Heru Handika, Anang S. Achmadi, Michael J. Polito, Kevin C. Rowe, Jacob A. Esselstyn

**Affiliations:** Florida Museum of Natural History, University of Florida, Gainesville, Florida, 32611, United States; Museum of Natural Science and Department of Biological Sciences, Louisiana State University, Baton Rouge, Louisiana, 70803, United States; Biological Sciences, Ohio University, Athens, Ohio, 45701, United States; Museum Zoologicum Bogoriense, Research Center for Ecology and Ethnobiology, National Research and Innovation Agency (BRIN), Cibinong, Jawa Barat 16911, Indonesia; Department of Oceanography and Coastal Sciences, Louisiana State University, Baton Rouge, Louisiana, 70803, United States; Department of Ocean Sciences, University of California, Santa Cruz, California, 95064, United States; Sciences Department, Museums Victoria, Melbourne, Victoria 3001, Australia

**Keywords:** Community Niche Space, Functional Morphology, Stable Isotopes, Murinae, Bayesian, Sulawesi, Indonesia

## Abstract

Communities that occupy similar environments but vary in the richness of closely related species can illuminate how functional variation and species richness interact to fill ecological space in the absence of abiotic filtering, though this has yet to be explored on an oceanic island where the processes of community assembly may differ from continental settings. In discrete montane communities on the island of Sulawesi, local murine rodent (rats and mice) richness ranges from 7 to 23 species. We measured 17 morphological, ecological, and isotopic traits, both individually and grouped into 5 multivariate traits in 40 species, to test for the expansion or packing of functional space among nine murine communities. We employed a novel probabilistic approach for integrating intraspecific and community-level trait variance into functional richness. Trait-specific and phylogenetic diversity patterns indicate dynamic community assembly due to variable niche expansion and packing on multiple niche axes. Locomotion and covarying traits such as tail length emerged as a fundamental axis of ecological variation, expanding functional space and enabling the niche packing of other traits such as diet and body size. Though trait divergence often explains functional diversity in island communities, we found that phylogenetic diversity facilitates functional space expansion in some conserved traits such as cranial shape, while more labile traits are overdispersed both within and between island clades, suggesting a role of niche complementarity. Our results evoke interspecific interactions, differences in trait lability, and the independent evolutionary trajectories of each of Sulawesi’s 6 murine clades as central to generating the exceptional functional diversity and species richness in this exceptional, insular radiation.

## Introduction

Whether at a local, continental, or global scale, species richness is not evenly distributed across the landscape. This unevenness emerges from both environmental and resource heterogeneity among communities, and from the interactions among the species within local communities.

Local species richness is often positively correlated with the complexity of habitat structure and diversity of available resources (Tews et al. 2004). The observation that different localities with similar habitat structure, resource availability, and historical access often contain ecologically similar communities led to the prediction that species only co-occur if they partition niche space along some axis (herein we consider the niche to be the size and shape of multivariate ecological space that a species utilizes), otherwise one will be excluded through competition (Hutchinson 1957, MacArthur & Levins 1967, May & MacArthur 1972, Brown & Lieberman 1973, Brown 1975, Pianka 1974, M’Closkey 1978). Competitive exclusion can be mitigated if two co-occurring species use a narrower breadth of resources, producing a more densely packed community niche space (“niche packing”), or if they exploit habitats or resources that are unused or non-existent in low-resource or species-poor communities, leading to a larger community niche space (“niche expansion”; MacArthur 1965, 1970, Pigot et al. 2016, Oliveira et al. 2020). The foundational work on competition’s role in community assembly relied on empirical data from continental communities with equal biogeographic accessibility but variation in primary productivity or habitat complexity (Brown & Lieberman 1973, Brown 1975, M’Closkey 1978, Karr & James 1975, Pianka 1974, Weiher & Keddy 1995), or from archipelagos where island size or geographic complexity determines resource availability and habitat area, both of which influence overall species richness (Wilson 1961, Diamond 1975, Lister 1976, Gillespie 2004, Losos 2009, Losos & Ricklefs 2009). Less explored are insular areas of similar habitat structure and resource availability, but with discrete communities that vary in species richness (“species richness anomalies”, Swenson et al. 2016 pg. E83). Yet, these anomalies offer powerful systems for interrogating the role of competition in the distribution of functional diversity within and among communities because the effects of abiotic processes such as habitat filtering are minimized (Swenson et al 2016, Li et al. 2017). Most studies of species richness anomalies have examined plants in continental settings (Latham & Ricklefs 1993, Swenson et al. 2016, Xu et al. 2019) where environmental and historical processes influence the regional species pool. Few, if any, studies have tested the role of competition in the assembly of discrete communities nested within an insular setting where species pools are formed through long-distance colonization and in-situ diversification.

Oceanic islands are often hotspots of diversification and endemism, and their unique biogeographic and historical conditions can help illuminate the ecological and evolutionary processes that structure local communities (Losos & Ricklefs 2009). Lineages that arrive and diversify in isolated locations regularly undergo ecological shifts that result in behaviors or phenotypes that are uncommon or non-existent in source communities (Carlquist 1966, Millen 2006, Pinto et al. 2008, Esselstyn et al. 2012, 2015, 2021, Stroud & Losos 2016). The presence of high functional and ecological disparity within an endemic radiation may affect community structure in multiple ways. First, there is some evidence that community assembly occurs differently on continents and islands. Using an assortment of phenotypic and behavioral proxies for resource use across a variety of spatial scales, niche packing is frequently invoked as the primary process of assembly in species-rich animal communities on continents (Brown 1975, Karr & James 1975, M’Closkey 1978, MacArthur & MacArthur 1961, Pianka 1974, Pigot et al. 2016, Van de Perre et al. 2020). However, niche expansion has found some support among isolated island communities of many closely related species (Lister 1976). Second, multiple lineages evolving in sympatry within an insular setting often promotes niche divergence through adaptive diversification, reducing phylogenetic niche conservatism (Losos et al. 2003). As a result, the niche breadth of a community within an insular setting, despite the lower phylogenetic diversity, may equal that of a similar continental community. Importantly, quantifying trait differences following adaptive diversification among closely related species can illuminate which traits are most evolutionary labile and/or important for resource partitioning (Losos et al. 2003, Hiller et al. 2019, Dorey et al. 2020, Stroud 2021).

The murine rodent fauna (rats and mice in the subfamily Murinae) of Sulawesi, Indonesia, a mountainous, wet tropical, oceanic island at the center of the Wallacean biodiversity hotspot (Figure 1), is an intriguing system for testing patterns of community niche occupancy.

**Figure 1:**
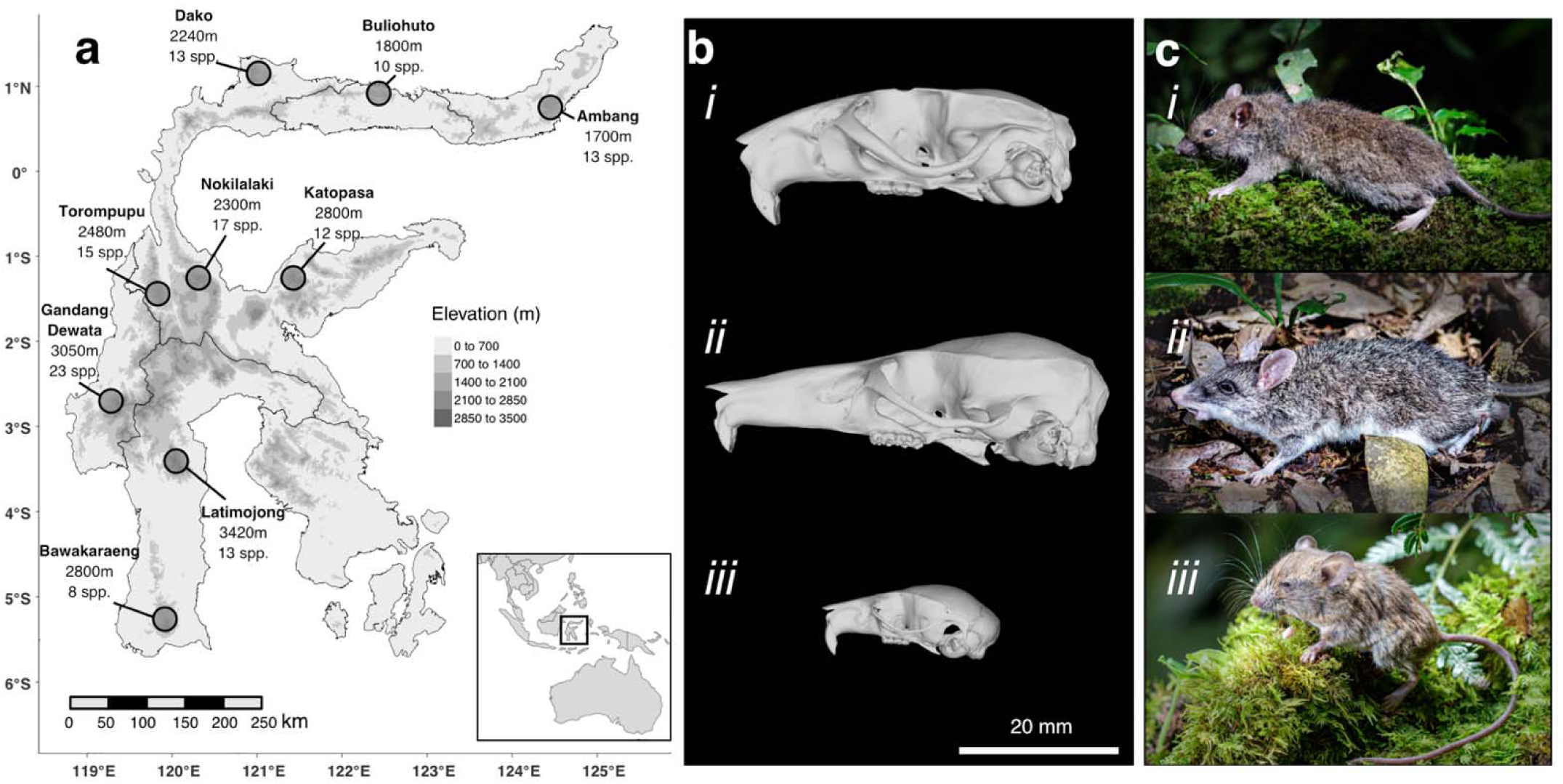
Small mammal surveys of nine mountains a) on the oceanic island of Sulawesi revealed varying murine rodent species richness across the island. Each mountain on the map is labeled with the maximum elevation and the number of murine rodent species present. All but Latimojong were surveyed within 600m of the summit. b) The diversity of Sulawesi murines is exceptional and includes unique forms that live alongside species with more “typical” ecologies and morphologies, as demonstrated from surfaces of cranial μCT scans of *i* – *Rattus hoffmanni,* a “typical” murine morphology and ecology, *ii* – the shrew rat *Echiothrix leucura* with its extremely elongate rostrum and soft invertebrate diet, *iii* – the arboreal *Haeromys minahassae*, with a short rostrum and very small size. c) Photographs: *i* – *Rattus hoffmanni*, *ii* –*Echiothrix leucura*, and *iii* – *Haeromys minahassae*.

First, the murine diversity of Sulawesi is exceptional, with at least 48 endemic species stemming from seven colonist ancestors that arrived from circa 6 Ma to < 1 Ma (Figure S1, Rowe et al. 2019, Handika et al. 2021). While some colonists spawned small radiations of species, others are evidenced by only a single living species (Table 1, Figure S1). Each clade is widespread on Sulawesi and most contribute to species richness of all local murine communities on the island. Second, Sulawesi contains some of the most unusual rodent forms found anywhere (Figure 1b, Esselstyn et al. 2012, 2015, Rowe et al. 2014) living in sympatry with more typical “rat-like” morphologies and ecologies. Sulawesi murines have an array of cranial shapes that reflect their dietary preferences (Figure 1b; Esselstyn et al. 2012, 2015, Martinez et al. 2018), consume a broad range of foods such as fruit, seeds, fungi, leaves, roots, and earthworms (Musser 2014, Rowe et al. 2016a), have body sizes ranging from 10 to 500g, and occupy a variety of locomotor modes (arboreal; scansorial; terrestrial; amphibious, Nations et al. 2021). Third, due to the topographic complexity of Sulawesi and the positive correlation between elevation and small mammal diversity in this region (Heaney 2001, Esselstyn et al. 2021), the murine communities on the islands are partitioned into discrete, montane assemblages (Figure 1). The local montane murine communities, defined as the species living on a mountain from the upper-lowland forest to the peak, range from 7 to 23 species, with the upper limit being, to our knowledge, the most diverse local community of closely related terrestrial mammals on Earth. Lastly, the variation in local community richness does not appear to be determined by environmental or habitat differences among mountains but is likely the result of the colonization process outward from the central core to the peninsulas during the island’s formation (Hall 2013, Nugraha & Hall 2018, Handika et al. 2021). Maximum elevation, which may correlate with the area occupied by different habitats, appears to play only a minor role in species richness (Figure 1a), and net primary productivity is nearly constant across the island’s montane regions (Imhoff et al. 2004), suggesting that environmental filtering, often a central process of community assembly (Webb et al. 2002, Cavender-Bares et al. 2004, Li et al. 2017), does not greatly affect the functional diversity of Sulawesi’s murine communities. Together, these properties generate a fascinating hierarchical organization: a species pool of small mammals with disparate ecologies and morphologies living in discrete montane communities that, due to their habitat similarity and striking disparity in richness, represent species-richness anomalies, all of which lie within a remote, oceanic island. Such a system presents a compelling natural experiment for testing alternative hypotheses of how communities assemble to occupy ecological niche space

**Table 1:** Ages of six clades descended from Sulawesi colonists. Median age in millions of years is reported with 95% credible intervals in parentheses. Species sampled reports the number of species in this study with overall clade richness in parentheses and demonstrates near complete sampling. The two unsampled *Margaretamys* species were not detected in the localities in this study. All ages taken from Rowe et al. 2019. *There were likely two colonizations by the ancestors of native *Rattus* spp., the second of which occurred 1.16-0.6 Ma (Rowe et al. 2019), however, all *Rattus* in this study form a clade relative to other Sulawesi murines. Two human commensal *Rattus* spp. found on Sulawesi were excluded. ^†^The age of arrival of the Haeromys clade is unknown as other species of Haeromys from Borneo have yet to be included in phylogenetic analyses.

| Clade | Crown Age | Species Sampled |
| --- | --- | --- |
| Echiothrix | 5.11 (4.49-5.69) | 10 (10) |
| Maxomys | 3.72 (3.21-4.29) | 5 (5) |
| Bunomys | 3.45 (3.09-3.81) | 15 (15) |
| Margaretamys | 2.92 (2.29-3.51) | 2 (4) |
| Rattus* | 1.57 (1.22-1.95) | 6 (6) |
| Haeromys<sup>†</sup> | Indeterminate | 1 (1) |

**Table 1:** Ages of six clades descended from Sulawesi colonists.
| Clade | Crown Age | Species Sampled |
| --- | --- | --- |
| Echiothrix | 5.11 (4.49-5.69) | 10 (10) |
| Maxomys | 3.72 (3.21-4.29) | 5 (5) |
| Bunomys | 3.45 (3.09-3.81) | 15 (15) |
| Margaretamys | 2.92 (2.29-3.51) | 2 (4) |
| Rattus* | 1.57 (1.22-1.95) | 6 (6) |
| Haeromys† | Indeterminate | 1 (1) |

Here, we test whether increased species richness in environmentally similar communities of montane rodents leads to the expansion or packing of ecological niche space. Observational data, including behavior, life history, and movement patterns, are scarce for the nocturnal, secretive Sulawesi murines. Much of what we know about these species comes from museum specimens and their associated metadata (e.g., locality and habitat details, forest strata preferences, and morphological measurements). Therefore, to infer the ecological niche breadth of each montane assemblage, we quantify diet, trophic dimension, and microhabitat using 12 individual functional and ecological traits and five multivariate trait complexes to estimate both the volume and density of community functional space, which we define as the sum of the n-dimensional functional spaces of the species therein. Importantly, individual traits or groups of traits related to the same niche axis can reveal distinct, interacting processes (Spasojevic & Suding 2012, Pigot et al. 2016, Kohli et al. 2021). Additionally, we examine the role of phylogenetic niche conservatism and trait lability in the assembly of these nine montane communities by testing for the influence of phylogenetic diversity and inter-clade variation on functional space occupancy.

## Materials and Methods

### The distribution of species-

We compiled occurrence records from nine small mammal inventories of mountain regions on Sulawesi, Indonesia, including one well-documented mountain surveyed from 1973 to 1976 by Guy Musser and colleagues (Mt. Nokilalaki; Musser, 2014) and eight mountains surveyed between 2011 and 2016 (Ambang, Bawakaraeng, Buliohuto, Dako, Gandang Dewata, Katopasa, Latimojong, and Torompupu; Figure 1a).

All surveys began in lower primary forest near the line of anthropogenic forest clearing (1100m to 1500m) and extended to upper-montane forests. All surveys extended to within 600m elevation of the summit except for Latimojong (highest survey site at 2535m, summit at 3400m). Trapping records show that there are no Sulawesi murines restricted to elevations above 2500m, or to habitats within 600m of the summit (Musser 2014). Historical surveys by Musser lasted several months and were conducted over four years, employing a mix of snap traps and live traps. Modern surveys (2011-16) lasted an average of 17 days (11-25) and employed similar collection methods, including a mix of snap traps, live traps, and 20-30L pitfall buckets. All the murine rodent species known from the sampled localities (Musser 2014, Wilson et al. 2019) were collected during these modern surveys, indicating a thorough sampling effort. Five new taxa that were discovered during these expeditions have been described (Musser 2014, Esselstyn et al. 2012, Rowe et al. 2014, Esselstyn et al. 2015, Rowe et al. 2016b) and several new locality records resulted (Achmadi, et al., 2014; Handika, et al., 2021). Specimens from all surveys were deposited in the Museum Zoologicum Bogoriense (MZB), Bogor, Indonesia; the American Museum of Natural History (AMNH), New York, USA; Museums Victoria (MV), Melbourne, Australia; the Museum of Vertebrate Zoology (MVZ), Berkeley, USA; the Field Museum of Natural History (FMNH), Chicago, USA; and the Louisiana State University Museum of Natural Science (LSUMZ), Baton Rouge, USA.

### Functional trait data collection and processing-

We compiled or generated functional trait values for 11 continuous traits and one discrete trait for all available Sulawesi murines from each of the nine communities. Individual Sulawesi murine species exhibit little intraspecific morphological variation among localities, far less than the morphological differences among species (Musser 2014), and we therefore combined measurements of individual species from multiple localities and estimated trait distributions using probabilistic methods to overcome the limited availability of some traits. Detailed information on traits, sample sizes, data processing, and multivariate trait composition are available in Table S1. All data, models, and output files are available in Dryad Repository (to be added prior to publication).

### Morphological data collection-

We assembled external measurement data from 630 specimens, including head-body length (mm), tail length (mm), hind-foot length (mm), ear length (mm), and mass (g), from previously published sources (Wilson et al. 2019, Nations et al. 2021) and online museum databases. Measured specimens were from the nine surveyed mountains and other localities on Sulawesi. To obtain ecologically relevant features of external measurements and mitigate the influence of size in some of our analyses, we calculated three commonly used ratios: Relative tail length (tail length / (head-body length + tail length)), relative hind-foot length (hind-foot length / head-body length), and relative ear length (ear length / head-body length) (Nations et al. 2021, Table S1).

The shape of a rodent’s skull and lower jaw provides a wealth of indirect ecological information on foraging, feeding, and sensory processing, and is often used as a proxy for fundamental dietary niche (Samuels 2009). We generated μCT scans of the cranium of 64 specimens from 38 species and the mandible of 61 specimens from 36 species (Table S1). Scans were generated from specimens collected in the nine surveys, as well as from previously collected museum materials. Stacks of 2D Tiff files were imported to MorphoDig, where 3D landmarks were placed on cropped volume renderings (Lebrun, 2018). We placed 67 cranial landmarks (Figure S2) on the left side of the skull, unless damage caused us to use the right side, in which case we reversed the rendering on the Z-axis. In separate renderings, we placed 20 landmarks on the left dentary of the mandible (Figure S2). Landmarks were exported from MorphoDig as .stv files and imported into the R package geomorph v.4.0.3 for processing (Adams et al. 2021, Baken et al. 2021). Missing landmarks (11 of 4288 cranial and three of 1220 mandible landmarks) were imputed, a generalized Procrustes analysis (GPA) superimposition was performed, and shape coordinates were subjected to a principal components analysis. We retained the centroid size (an estimate of total size) and the scores from the first 36 principal components of the cranium and 20 axes of the mandible, each representing >95% of the shape variation of the element.

### Stable isotope data collection-

Approximately 1-2 grams of hairs were plucked from the rump of 286 dry museum specimens collected on six focal surveys (Ambang, Bawakaraeng, Buliohuto, Dako, Gandang Dewata, and Latimojong). Isotopic values can vary regionally (Fry 2006); therefore, we collected hair samples from multiple individuals of each species from each locality (mean = 4.6 specimens/species/locality, range = 1-14). Nitrogen stable isotope values (δ^15^N) act as a proxy for consumer trophic position as they generally increase by 3–5% per trophic level (DeNiro & Epstein 1981). Carbon isotope values (δ^13^C) generally exhibit little to no change with trophic position and are commonly used as proxies of consumer’s basal carbon resource use (e.g. use of differing primary production energy pathways; DeNiro & Epstein 1978). Combined, these two metrics are commonly used to quantify the “isotopic niche” of consumers, which can act as a useful proxy of species realized dietary niche (Newsome et al. 2007, Ben-David & Flaherty 2012). Stable isotope values are reported in delta notation in per mil units. Samples were processed at the Stable Isotope Ecology Laboratory, Department of Oceanography & Coastal Sciences, Louisiana State University. Details on sample cleaning, processing, and analysis are in the Supporting Methods.

### Locomotor mode data collection-

We used the locomotor classification scheme from Nations et al. (2019, 2021) to group each Sulawesi murine into one of four discrete locomotor modes: Arboreal, (climbing is integral to survival); General (navigates a variety of substrates and habitat strata); Terrestrial (on the ground surface); and Amphibious, (dependent on aquatic habitats for foraging).

### Combined traits-

We combined our 11 continuous traits into five multivariate traits that represent distinct niche dimensions: (1) head shape (cranium shape PC 1-36 and mandible shape PC 1-20); (2) isotopic niche space (δ^15^N and δ^13^C stable isotope values); (3) body proportions (head-body length, relative tail length, relative hind-foot length, and relative ear length); (4) body size (log(mass) and head-body length); and (5) total morphological shape (all nine morphological traits). We did this by combining the predicted trait values for each community (see below).

### Estimating species’ trait values-

The sample size of morphological and isotopic data varied between species and mountain communities, and in some cases was limited to one individual, such as with the amphibious *Waiomys mammasae*, a species known from a single specimen (Rowe et al. 2014). To mitigate uneven sampling, we estimated a probability distribution of species trait values using partial-pooling in a multilevel Bayesian model. Unlike complete pooling (one global mean estimated for all combined samples) or no-pooling (one mean estimated per species, independent of all others), partial-pooling estimates a mean for each species as well as the variance among species, which serves as an adaptive prior that is common to all the species’ means (McElreath 2020). The Bayesian partial-pooling modeling allowed us to incorporate intraspecific variation in trait value predictions while avoiding point estimates such as averages, which discard valuable information. This approach prevents unbalanced estimates by using the trait-variance probability estimates from well-sampled species to inform variance estimates of species with fewer samples (Gelman & Hill 2006, McElreath 2020). All analyses were conducted in the probabilistic programming language Stan (Carpenter et al. 2017) within the R library brms v. 2.17.0 (Bürkner 2018). Subsequent data processing and figuring relied on the R libraries tidyverse v. 1.3.1 (Wickham et al. 2020), furrr v. 0.3.0 (Vaughan 2021), and tidybayes v. 3.0.2 (Kay 2020). All data, scripts, and output files are available in Zenodo Repository (to be added prior to publication) and on GitHub (to be added).

For each continuous trait (Table S1), we used the trait value as the response variable, and used species as a group-level predictor. All traits were scaled to unity prior to analyses. We used the student-*t* distribution to describe the response variable to minimize the influence of rare, extreme observations (a.k.a. ‘robust regression’, Kruschke 2013, McElreath 2020). Each model included four chains with 4000 iterations of warm-up and 1000 sampling iterations. Posterior predictions from the four chains were combined, resulting in 4000 samples per trait per species. To mitigate the potential geographic signal in stable isotope values (Fry 2006), and facilitate randomization (see below), the posterior estimates of the δ^15^N and δ^13^C stable isotope values were estimated with models that included an additional ‘community’ group-level effect, but otherwise were identical to the models described above (Supporting Methods). These stable isotope models generated posterior values of species’ isotopic measurements conditioned on the locality, enabling comparisons among communities and randomized sampling for null models (Table S1, Figure S3). Full details of the model, prior, and chain-estimation are described in the Supporting Methods.

### Estimating functional space volume-

We defined functional space volume as the total volume of n-dimensional trait space occupied by species in a community, and for consistency and clarity we use the term ‘functional space volume’ for all individual and multivariate traits, regardless of dimension. We used the sum of the variance of each trait value to estimate the volume of community functional space for each community, which is less sensitive to outliers and outperforms other metrics such as ellipse volumes, convex hull volumes, and hypervolumes, especially for functional spaces with many axes (Li et al. 2017, Guillerme et al. 2020). We estimated a posterior distribution of the variance of each trait for each community by first grouping the species’ trait value estimates by community. Then, for each of the 4000 posterior draws (one draw ranging between 7 and 23 values, depending on the richness of the community), we estimated the variance of the trait, resulting in a distribution of 4000 variance values for each trait for each community. We estimated the combined multivariate trait space variance by summing the variances of each trait, then dividing by the number of traits. We estimated the 89% probability values for the variance of each individual and combined trait space. To estimate locomotor mode variance, we dummy coded locomotor mode into four binary columns (one per mode) and performed a redundancy analysis (Legendre & Legendre 2012) with the rda() function in the vegan package (Oksanen et al. 2019). We then extracted the PC scores for each species and calculated the sum of variances of the three PC axes for each community.

### Estimating functional space density-

We estimated the density of species within functional trait space with the mean nearest neighbor (NN) metric (Guillerme et al. 2020). Our methods follow the estimates of functional space volume above. First, we grouped species’ trait values by community, then estimated the individual trait density for each posterior draw using the NN metric in the disparity() function of the R library dispRity (Guillerme 2018), resulting in a distribution of 4000 NN values for each trait and combined trait space. To estimate locomotor mode density, performed a redundancy analysis on the dummy-coded locomotor data as above. We then extracted the PC scores for each species, grouped the species by community, and estimated the NN value for each community as with the continuous traits above, in this case generating only a single NN value per community rather than a distribution.

### Null models of functional space volume and density-

We used null models to determine the difference between the functional space of our nine communities and randomly assembled communities. For each community we created 1000 randomized null communities of *n_community_* species for each of the nine localities using the independent swap algorithm (Gotelli 2001). We then calculated both volume and density as above for each individual (n=12) and combined (n=5) trait for each of the 1000 randomized community samples. Standardized effect size (SES) was calculated as:

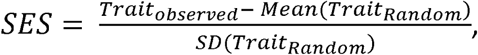

where *Trait_observed_* is the vector of 4000 samples of the density or volume of the given trait, Mean(*Trait_Random_*) and SD(*Trait_Random_*) are the mean and standard deviation of the density or volume values of the given trait from the 1000 random species assemblies. Positive SES values indicate greater than random functional space volume or lower than random functional space density (overdispersion), meaning that species in that community are occupying functional space outside the range of other communities and/or their niches are farther apart in functional space (niche expansion). Negative SES values indicate lower than random functional space volume and greater than random functional space density (underdispersion or clustering), meaning that species in that community are occupying less ecological space than other communities and/or their niches are closer together in functional space (niche packing; Oliveria et al. 2020).

### Effect of species richness on community functional space volume and density-

If niche expansion is the primary mode of resource partitioning in Sulawesi murine communities, then we expected that functional space volume will increase with increasing species richness, and that functional space density will remain stable as richness increases. Whereas if niche packing is occurring, we expected that functional space density will increase with species richness, and that there will be no effect of richness on functional space volume. To quantify the effect size of richness ((*β_richness_*) on functional space volume and density, we used Bayesian linear regression models that include the measurement error of the predictor variable to estimate the effect size of species richness on volume or density. We used species richness as the predictor variable, and the mean and standard error of the trait estimates for each community as the response. Measurement error models contain vastly more information per observation than a single point observation and provide robust estimates despite the small number of sampled communities (Bürkner 2018, McElreath 2020). We also calculated the Bayesian R^2^ value for each regression (Gelman et al. 2019). The models, priors, and chain estimations are detailed in the Supporting Methods.

### Phylogenetic diversity as a path to trait disparity-

The ecological space occupied by a community may depend on which lineages or clades are present (Webb 2000). Trait disparity appears to vary among clades of Sulawesi murines, and if so, then the volume and densities of functional spaces may be more influenced by phylogenetic diversity than ecological factors. To determine how phylogenetic diversity mediates functional space occupancy in the nine murine communities, we estimated the phylogenetic diversity (PD) of each community using Faith’s metric of branch length (Faith 1996). We removed all but the Sulawesi species (n = 35) from a time-calibrated phylogenetic hypothesis of Murinae (Nations et al. 2021). For the analyses described above, we had trait data for four species that are not included in this phylogeny: *Maxomys wattsi, Rattus bontanus*, *Rattus mollicomulus*, and *Rattus marmosurus*. We manually added these four species into the tree using the R package phytools (Revell 2012). Details are found in the Supplemental Methods. We used the R package picante (Kembell et al. 2010) to estimated Faith’s metric of PD, and to sample 1000 random communities in order to calculate the SES value of PD (SES PD). We used linear modeling in brms to estimate the effect size of species richness on SES PD. *Haeromys minahassae* is the only representative of its genus on Sulawesi, and, due to its large phylogenetic distance from other Sulawesi murines, may have an oversized impact on estimates of community PD. Therefore, we repeated the estimates of SES PD and effect size above without *Haeromys minahassae*. To quantify the trait disparity within and among Sulawesi murine clades, we grouped the posterior distributions of the species’ predicted trait values by clade, then estimated the variance of each of the 12 univariate traits. We plotted the trait value variance for each community along with the species’ predicted trait values (Figure S4)

## Results

### Models of trait values-

All parameters in the Bayesian multilevel model estimates of trait-value distributions and measurement-error estimates of the species richness effect on trait values had ESS > 1000 and a Gelman-Rubin diagnostic *R̂* ≤ 1.01, indicating convergence. Raw volume and density values differed from the volume and density of SES. To avoid the confounding influence of species richness in volume and density estimates we report and discuss only the SES values (Swenson 2014).

### Effects of Species Richness on Functional Space Volume and Density-

Functional space volume was positively correlated with species richness in seven of the 12 traits (locomotor mode, tail length, head-body length, cranium shape, mass, δ^15^N, and hind foot length), as shown by the positive (J_richness_ slope values from the Bayesian linear models (Figures 3, Table S3) and the greater variance with higher community richness (Figure 3). The remaining five traits (ear length, cranium size, dentary size, dentary shape, and δ^13^C) did not substantially increase in volume with increased species richness. Among the multivariate traits, the SES values of total morphological shape increased substantially with species richness while head shape and isotopic niche space show a moderate probability of increasing with species richness (Figures 3, 4, Table S2, S3).

**Figure 2:**
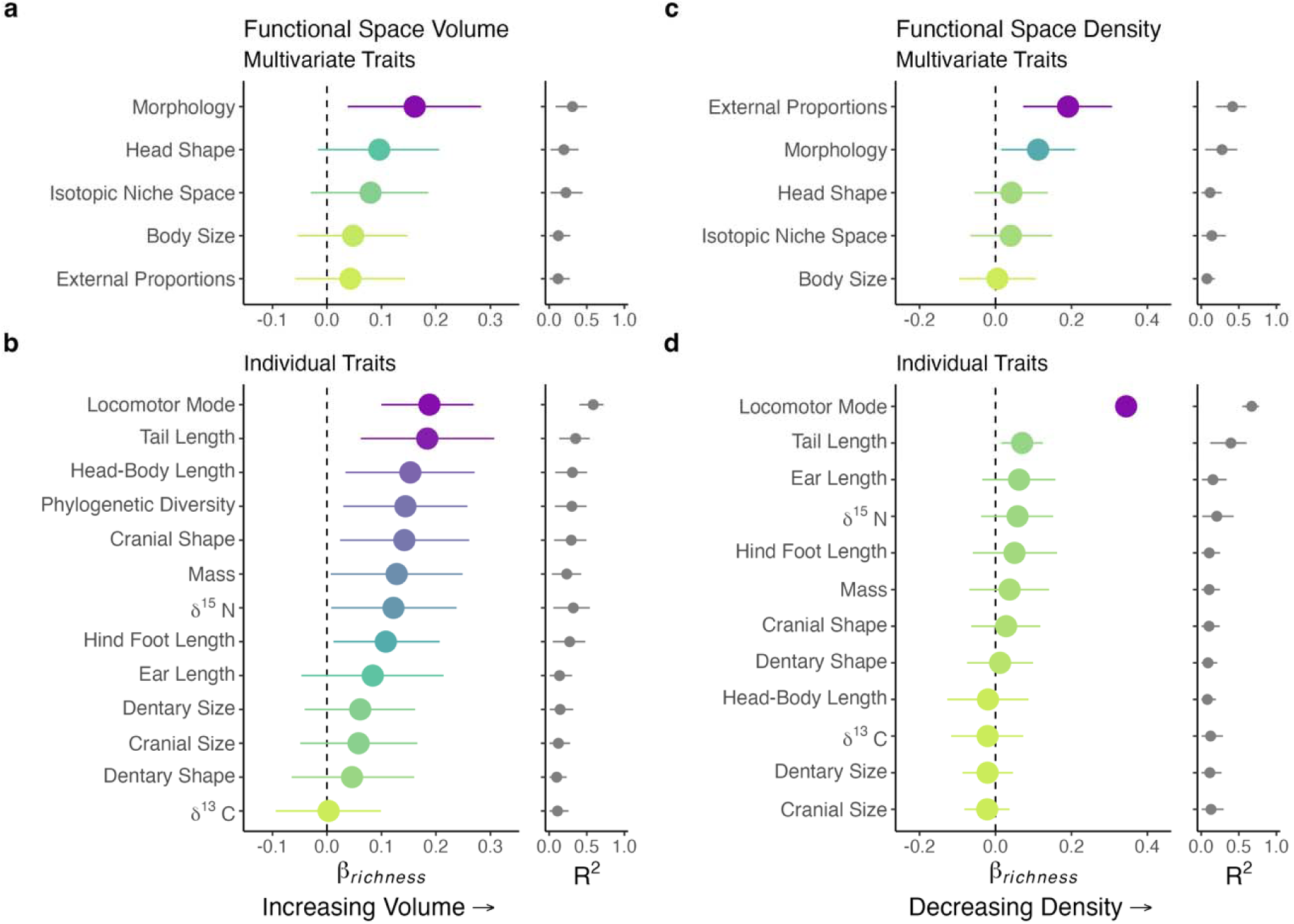
The effect of species richness on community functional space volume (a, b) and density (c, d). X-axes show the (*β_richness_* estimates (the regression slope) for each trait space (y-axis) on the left and the Bayesian R^2^ for each (*β_richness_* estimate on the right. Colored point intervals show 89% probability of (*β_richness_* estimates (effect size), with color varying by (*β_richness_* value. Black point intervals show 50% posterior estimates of Bayesian R^2^ for each trait space. Density was estimated using the mean nearest neighbor (NN), and a high NN distance indicates low density. Four of the five multivariate traits (**a**) and seven of the 12 individual traits (**b**), show an increase in functional space volume (trait variance) with greater richness (i.e., positive (*β_richness_*). Phylogenetic diversity also increases with species richness (**b**). All multivariate traits (**c**) and individual traits (**d**) show a stable or, surprisingly, decreasing functional space density (increased NN distance) with greater species richness.

**Figure 3:**
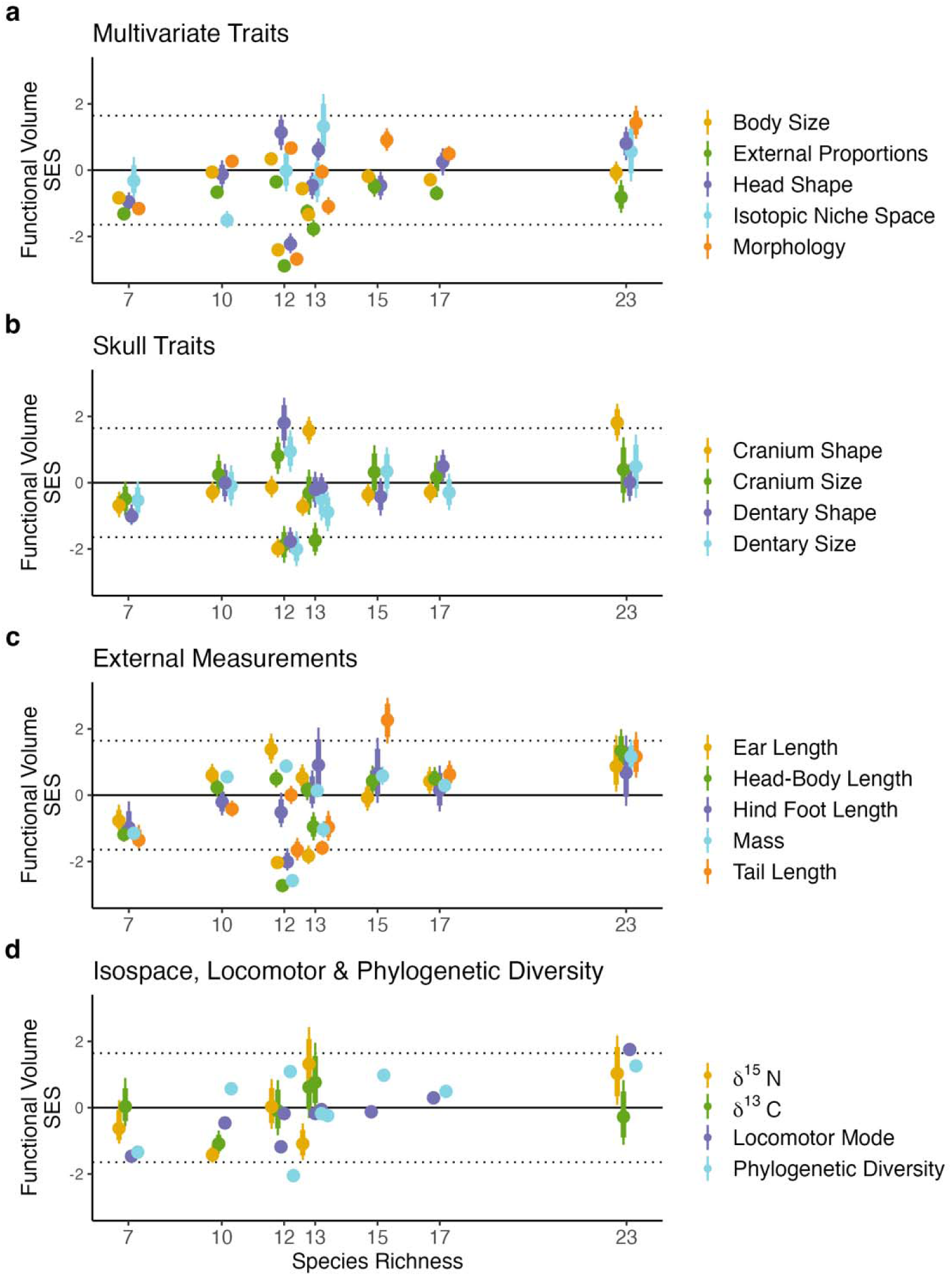
Estimated functional volume SES plotted against species richness: **a**) Multivariate trait volumes, **b**) Skull traits, **c**) External measurements, and **d**) Isotopic measures, locomotor mode, and phylogenetic diversity. Points represent the mean SES values and error bars indicate the 89% credible intervals. X-axis tick marks show species richness for each community. Values equal to zero are consistent with null expectations, positive values indicate overdispersion, and negative values show underdispersion (trait clustering). Dotted lines depict the 89% interval of the null distribution. The plot of trait densities is shown in the supporting information (Figure S4).

Linear regression showed that species richness had little to no effect on functional space density (NN SES values) for 11 of the 12 individual traits and three of five multivariate traits (Figures 3, S4, Table S2), consistent with niche expansion. The exceptions were a positive (*β_richness_* slope (i.e., decreased density with richness) for locomotor mode, total morphological shape, and external proportions, suggesting an extreme overdispersion of morphology and locomotor diversity in the richest communities.

### Phylogenetic Diversity and Functional space-

Phylogenetic diversity (PD) increased with species richness (Figures 3b, 4d, Table S2). Including *Haeromys minahassae* in the estimates of PD changed the PD SES values of the individual communities but had minimal impact on linear regressions (Table S4). Results including *H. minahassae* had a slope ((*β_richness_*) of 0.143 (0.03, 0.252) while excluding *H. minahassae* generated a slope of 0.169 (89% C.I. of 0.041, 0.295), and we therefore present the results that include *H. minahassae*. Katopasa, a community of 12 murines on Sulawesi’s Eastern Peninsula, has the lowest PD SES value followed by Bawakaraeng on the Southwestern Peninsula (seven species). Gandang Dewata, the richest community (23 species), has the highest PD SES value (Figure 3a, Table S2). As suspected, functional space volumes vary between clades, though not consistently (Figure S5). For example, the Echiothrix clade has the highest variance in cranial shape, dentary shape, and dentary size but low trait variance for hind-foot length and δ^15^N. In contrast, the Maxomys clade has low variance for all trait values except tail length and δ^15^N (Figure S5). This results in different densities of functional space occupation among traits, where some trait volumes, such as cranial space, are strongly influenced by phylogeny, while others, such as isotopic niche space, have high and low values distributed among clades (Figure 4).

**Figure 4:**
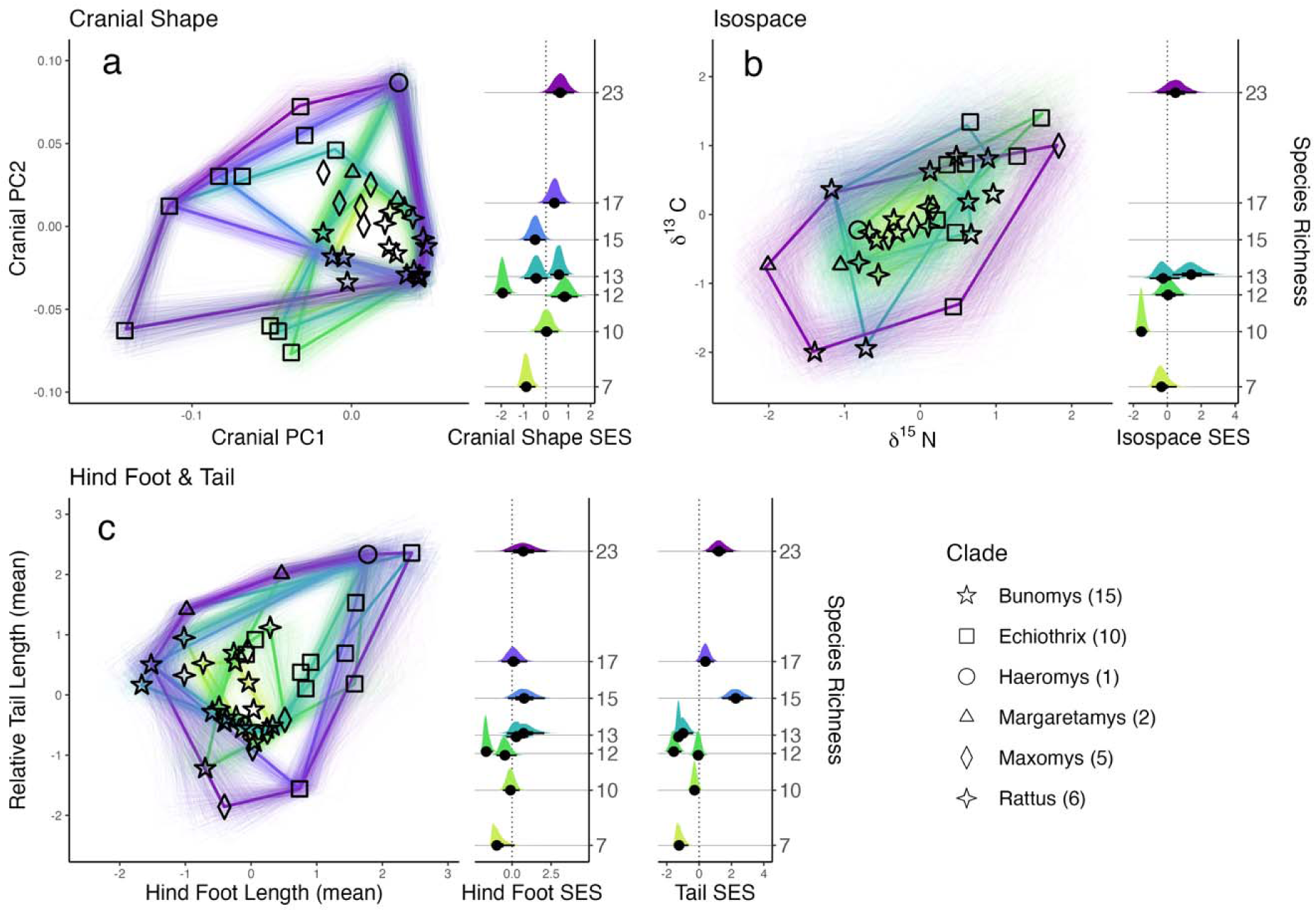
The mode of functional space filling varies among traits. The bivariate plots on the left depict the 2D functional spaces. Each black shape is the mean value of a particular species in each clade (shown in legend). The convex hull colors match the color of the community functional volume SES distributions in the right-hand columns. The thick convex hull lines are mean values, and 500 random samples from the posterior of each trait are shown in thin hull lines. The number of species in each community and each clade used in this study is shown next to the SES distributions in parentheses. All values were scaled to z-scores prior to analyses. **a**) Cranial morphospace values are mostly clustered tightly near the mean of each axis, apart from Echiothrix species and the single *Haeromys* species. Intraspecific variance is also relatively low on these axes. **b**) Intraspecific variance is high in isotopic niche space. High and low δ^15^N values are distributed among clades, but that is not the case for δ^13^C values. **c**) Large values of hind foot length belong to members of two clades, while large and small tail length values are dispersed among all six clades, reducing the influence of phylogenetic diversity on tail length disparity.

## Discussion

Variation in species richness among communities of closely related species that inhabit similar environments provides a unique window to explore how interspecific competition may affect community functional richness in the absence of confounding factors like environmental variation (Roughgarden 1976, Swenson et al. 2016). Unfortunately, these species richness anomalies are uncommon (Swenson et al. 2016, Van de Perre et al. 2020). Here we used a large dataset of ecological and morphological traits to estimate the changes in community niche occupancy across a richness gradient on an oceanic island. Though the trait estimates of some species necessarily stem from a small number of specimens, and therefore may be subject to error, our novel probabilistic approach incorporates measurement error and species-level variation into the posterior distribution and propagates this uncertainty through the estimation of community niche space. The functional space of most traits increased with greater species richness among the nine murine communities studied while there was no change in functional space density, consistent with limiting similarity. Locomotor mode disparity has the strongest positive relationship with species richness, but the functional volumes of skull and body shape (indicators of diet, locomotion, and microhabitat use in murines and vertebrates in general; Peters 1986, Losos 2009, Martinez et al. 2018, Nations et al. 2021), also strongly increase with species richness (Figure 2). Measurements of overall body size, cranial size, dentary shape and size, ear length, and δ^13^C values all demonstrate either a weak signal of niche packing, or no signal across the species richness gradient. While the volume of many functional spaces increases with richness, functional space density shows little correlation with species richness for most traits, and it surprisingly decreases with increased richness for external morphology and locomotor mode. These results, along with a general underdispersion of trait values in low-richness communities and overdispersion in high-richness communities (Table S2, Figures 4 & S4), suggest that species in rich communities mitigate competitive interactions by occupying underused niche space. We also found that increasing phylogenetic diversity is a means of increasing functional space occupation. Combined, these results point to species interactions as a mechanism for both phenotypic and phylogenetic overdispersion (Webb et al. 2002, Cavender-Bares et al. 2004, Li et al. 2017) and suggest that, given adequate resources, structural complexity, and evolutionary time, lineages can evolve to occupy unique regions of ecospace, often far from the average trait value, which minimizes niche overlap and cultivates exceptional richness.

Whether in a single desert valley (Brown 1975), or across continental (Maestri & Patterson 2016, Kohli et al. 2022) and global latitudinal gradients (Karr & James 1975, Pellissier et al. 2018), structurally complex habitats are thought to foster higher species diversity. High plant diversity creates a more complex, vertically structured habitat matrix for other plant and animal species to occupy and has long been tied to higher animal richness (Hutchinson 1959, MacArthur & MacArthur 1961, Scheffers et al. 2013, Oliveira & Scheffers 2019). Our estimates of locomotor-mode occupancy clearly demonstrate that vertical habitat partitioning is critical to maintaining high species richness in Sulawesi murines (Figure 2, Table S2). Strikingly, the density of locomotor trait space decreases along the richness gradient, indicating a very high level of trait overdispersion (Figures 3c & S4). Arboreal, Terrestrial, and Amphibious locomotor modes each provide access to different microhabitats that contain similar resources. Among the 23 species found on Mt. Gandang Dewata, the amphibious *Waiomys mamasae*, the terrestrial *Paucidentomys vermidax*, and the arboreal *Sommeromys macrorhinos*, all consume invertebrates and have similar δ^15^N values, yet they are unlikely to compete for resources due to their distinct microhabitat use, a pattern observed in other insular communities of closely related vertebrates (e.g., Jamaican *Anolis*; Shoener 1974). It’s worth noting that when the lone amphibious species *Waiomys mamasae*, only known from Gandang Dewata, is removed from the data, the locomotor variance of Gandang Dewata remains the highest among the communities. Morphological measurements are often used to infer locomotor mode in a variety of eco-evolutionary contexts (Ricklefs & Travis 1980, Samuels and Van Valkenburgh 2009, Pianka et al. 2017, Verde-Arregoitia et al. 2019), and our estimates of tail-length variance, a trait correlated with locomotion in murines (Nations et al. 2021), increases with richness at nearly the same rate as locomotor mode (Figure 2). Importantly, our results suggest that locomotor mode rankings may be an effective way to estimate community locomotor variance where continuous trait data are lacking. Combining many traits into one multivariate measure of functional diversity is a common approach in evolutionary ecology. Ordination techniques were especially promoted to overcome pitfalls from early community ecology studies that used few, largely subjective measures of resource use (Ricklefs & Travis 1980). However, merging traits into multivariate axes masks trait-specific processes related to functional space (Spasojevic & Suding 2012, Astor et al. 2014). Indeed, our trait volume and density estimates reveal distinct patterns between individual and combined traits. For example, isotopic niche space, a combined signal of δ^15^N and δ^13^C values commonly used in terrestrial ecological studies, exhibits an equivocal signal (Figure 2b), but individual isotopic values reveal that the packing signal originates from the static δ^13^C value along the species richness gradient. The δ^15^N value, a signal of trophic level, expands along the richness gradient, a result that would be overlooked in multivariate analyses. In contrast to δ^15^N, dentary shape exhibits an equivocal signal, despite its presumed relationship to dietary niche (Maestri et al. 2016, Kohli et al. 2019). These opposing patterns are only illuminated by analyzing individual traits (Spasojevic & Suding 2012). If we performed this study using only cranial size and dentary shape, which are thought to capture the important axes of size and diet in murine rodents (Rowsey et al. 2019, 2020) and mammals in general (Prevosti et al. 2012, Grossnickle 2020), our results would suggest that functional volume does not increase with species richness.

The disparity of traits within a clade mediates the impact of phylogenetic diversity on community functional space volume. If trait values are phylogenetically clustered, then increasing phylogenetic diversity is necessary for community niche expansion to occur. But if trait values are phylogenetically overdispersed, niche packing, expansion, or, as we found, both could result from increased phylogenetic diversity. The distribution of traits among clades is of particular interest in communities that are assembled through a mix of colonization and in situ speciation, such as Sulawesi murines, Caribbean anoles, or Hawaiian spiders (Gillespie 2004, Losos 2009, Rowe et al. 2019). Niche divergence has been hypothesized to overcome niche conservatism in communities with an extended history of coevolution, such as those on oceanic islands, likely diminishing the signal of phylogenetic trait clustering (Losos et al. 2003). We find that niche divergence and conservatism may occur simultaneously on different functional traits. For example, the elongate, highly distinct skulls of some species in the Echiothrix clade, the descendants of the first murine colonists on Sulawesi, set them apart from other clades in skull shape and, despite their relatively low abundance, these species overcontribute to community cranial and dentary shape volumes (Figures 3b, 4b, S5). Unlike cranial shape however, the Echiothrix clade occupies a very constrained portion of δ^15^N trophic space. Additionally, trophic level estimates from δ^15^N values are notably dispersed among clades (Figures 4c, S5), and high phylogenetic diversity is not necessary for high isotopic niche space estimates. Are there innate differences in these traits that could lead to opposing patterns of niche conservatism?

The evolutionary lability of ecologically important traits determines the rate of convergence and divergence possible within a given time frame and can directly influence dispersion of trait values among species (Cavender-Bares et al. 2004). The traits with the highest within-clade variance — δ^15^N, tail length, body size — are all thought to be evolutionarily labile. Changes in tail length, body size, and intestinal tract morphology can occur on brief evolutionary time scales (Powell & King 1997, Naya et al. 2008, Kingsley et al. 2017, 2021), whereas morphological changes in cranial shape, such as substantial rostral elongation and the reduction of molar grinding area, may take much longer. Indeed, insular species are known to have rapidly expanded breadths of diet and labile morphological traits following colonization (Stuart et al. 2014) and subsequent speciation (Wilson 1959, 1961, Lister 1976, Millien 2006, Rowe et al. 2016a). But the extreme cranial morphologies of some Sulawesi murines, particularly those of the oldest radiation on the island (Echiothrix clade), are likely the result of a long process of in-situ evolution (Rowe et al. 2019). In other words, on shorter time scales, similarities in some slowly evolving traits may lead to higher divergences in more labile traits, as evidenced in the Maxomys and Bunomys clades (Figure S5). The opposite pattern may also occur, but only following sufficient evolutionary time. We propose that the evolutionary lability of traits is a determinant of trait value dispersion (Webb et al. 2002, Cavender-Bares et al. 2004), which directly relates to our inferences of functional space occupancy in Sulawesi murine communities. The theory of niche complementarity suggests that a pair of coexisting species that are similar in one trait should diverge in another trait (Schoener 1974). Yet, in an isolated setting, the lability of the traits in question necessarily determines the degree of complementarity possible within a given time frame. Linking trait lability with niche complementarity in an island system has important implications for the generation of functional and taxonomic diversity and may further illuminate the process of niche filling in insular, adaptive radiations.

## Conclusion

Here we provide evidence that limiting similarity in functional traits reflecting locomotion and microhabitat use plays an important role in the assembly of discrete, montane small mammal communities on an oceanic island. Our results contrast with recent studies that recovered niche packing as key to increased species richness in continental tropical vertebrate communities (Pigot et al. 2016, Peixoto et al. 201, Pellissier et al. 2018, Van de Perre 2020, Dehling et al. 2022, Hughes et al. 2022). Furthermore, our results counter the predictions that niche packing is expected to occur if resources remain constant among communities (Roughgarden 1976). Yet, Roughgarden (1976) posited that the predicted relationship between species richness and resources may be different on remote islands, as the distance from source populations may affect both community richness and the functional trait values of the species present. The organization of distinct montane communities on an oceanic island may be just such an example. The regional murine species pool of Sulawesi is itself structured by idiosyncratic immigration, diversification, and (presumably) extinction dynamics, and the resultant species’ functional traits. The species in these communities that occupy the edges of some functional spaces, such as cranial shape (Figure 4), represent morphologies and ecologies found only on Sulawesi or other large, oceanic islands. And the “imperfect isolation” (sensu Samonds et al 2013) of Sulawesi allows for other regional murine taxa, often represented by more “average” murine phenotypes, to periodically colonize the island, adding more species into the center of functional space (Figure 4). The presence of such disparate phenotypes and ecologies on Sulawesi (Esselstyn et al. 2012, 2015, Rowe et al. 2014) produces a larger functional space reserve than is available in most, if any, continental rodent systems, nurturing both niche expansion and high species richness. Therefore, the complex topography, isolation, abundant resources, and sequential colonization of Sulawesi might lead to species assembly processes that are typical of other large, oceanic islands, but are atypical of continental systems, “sky islands”, geologically younger islands, or regions with less abundant resources (deserts, high latitude habitats), but the presence of this general pattern remains untested.

