## Supporting Information for "The roles of isolation and interspecific interaction in generating the functional diversity of an insular mammal radiation" (from another version of this preprint: version 2, 2024-05-14)

### Supporting Methods

*Stable isotope sampling, processing, and analyses*- Approximately 1-2 grams of hairs were plucked from the rump of 286 dry museum specimens collected on six focal surveys (mean = 4.6 specimens/species/locality, range = 1-14). External contaminants and lipids were cleaned from the hair using a 2:1 chloroform: methanol solution (Hückstädt et al. 2012). Hair samples (0.6 ± 0.025 mg) were loaded in tin capsules and flash combusted on a Costech ECS 4010 elemental analyzer coupled to a Thermo-Fisher Delta Plus XP continuous-flow stable isotope ratio mass spectrometer. We normalized stable isotope values with the USGS-40 and USGS-41 glutamic acid reference materials. Stable isotope values, reported in delta notation (δ) in per mil units, were calculated with the following equation:

$$\delta X = \left[\left(\frac{R_{sample}}{R_{standard}}\right) - 1\right] x\ 1000$$

with X representing $^{13}C$ or $^{15}N$, and $R_{sample}$ and $R_{standard}$ representing the $^{13}C/^{12}C$ or $^{15}N/^{14}N$ ratios of sample and standard. Sample precision based on replicate reference materials was 0.1‰ for both $\delta^{15}N$ and $\delta^{13}C$ values. All samples were cleaned and processed by J.A.N. at the Stable Isotope Ecology Laboratory at Louisiana State University Department of Oceanography & Coastal Sciences.

*Estimating the partially-pooled distributions of trait values*- Posterior probability distributions for all continuous traits (cranium PC 1-36 scores, and mandible PC 1-20 scores, $\delta^{15}N$ and $\delta^{13}C$ stable isotope values, and external measurements) were estimated for each species using the same model and series of steps. First, trait values were scaled to unit variance (mean = 0, sd = 1), which placed all measurements on the same scale and informed our prior predictions (McElreath 2020). We used robust regression with the student-t distribution to minimize the influence of rare outlying response values (Kruschke 2013, McElreath 2020). The posterior distribution of each trait value for each species was estimated as:

$$
\begin{gathered}
y_i = Student(\upsilon, \mu_i, \sigma_i) \\
\mu_i = \alpha_{species[i]} \\
\alpha_j = Normal\ (\bar{\alpha}, \sigma) \\
\bar{\alpha} = Normal(0, 1) \\
\sigma = HalfNormal(0, 1) \\
\upsilon = 3
\end{gathered}
$$

where $y_i$ is the outcome (scaled trait value of each sample), $\alpha_{species[i]}$ is the unique value for each species, $\alpha_j$ is the multilevel prior distribution for all species 1…j , $\bar{\alpha}$ is the hyperparameter of the average trait value, representing the variation among species, and $\sigma$ the hyperparameter of the average species error. The tail-shape parameter of the student-t distribution, $\upsilon$, was set to three to minimize the influence of rare, extreme observations. Each model included four chains run for 5000 iterations, with 4000 warm-up and 1000 sampling iterations, resulting in a distribution of 4000 draws for each trait for each species. Convergence was suggested by an effective sample size (ESS) > 1000 and a Gelman-Rubin diagnostic $\hat{R} \leq 1.01$. To mitigate the geographic signal in stable isotope values (Fry 2006), and facilitate randomization (see below), we ran two additional models using $\delta^{15}N$ and $\delta^{13}C$ stable isotope values as responses, 'species' as one group-level predictor, as above, and included 'community' as a second group-level predictor. This model estimates two adaptive priors: one for all species and one for all communities. The model was:

$$y_i = Student(\upsilon, \mu_i, \sigma_i)$$
$$\mu_i = \alpha_{species[i]} + \gamma_{community[i]}$$
$$\alpha_j = Normal\ (\bar{\alpha}, \sigma)$$
$$\gamma_j = Normal\ (\bar{\gamma}, \sigma)$$
$$\bar{\alpha} = Normal(0, 1)$$
$$\bar{\gamma} = Normal(0,1)$$
$$\sigma = HalfNormal(0, 1)$$
$$\upsilon = 3$$

where $\gamma_{community[i]}$ is the is the unique value for each community, $\gamma_j$ is the multilevel prior distribution for each communities 1…j , $\bar{\gamma}$ is the hyperparameter of the average community isotope value. The remainder of the parameters remain the same as the equation above. This resulted in posterior values of species' isotopic measurements, conditioned on the community, enabling comparisons among communities and randomized sampling for null models (Table S1, Figure S3).

*The effect of species richness on community ecospace*- To quantify the effect of richness on community trait volume and density, we used Bayesian linear models that include the measurement error of the predictor variable to estimate the effect size of species richness on volume or density. Measurement error models contain vastly more information per observation than a single point observation and provide robust estimates despite the small number of sampled communities (McElreath 2020, Bürkner 2018). Our models took the form:

$$y_i \sim Normal(\mu_i, \sigma_i)$$
$$\mu_i = \alpha + \beta M_{true\ i}$$
$$M_{obs,i} \sim Normal(M_{true,i}, M_{SE,i})$$
$$M_{true,i} \sim Normal(0,1)$$
$$\alpha \sim Normal\ (0,1)$$
$$\beta \sim Normal(0, 1)$$
$$\sigma \sim HalfNormal(0, 1)$$

where $y_i$ is the species richness of each community, $M_{obs,i}$ and $M_{SE,i}$ are the observed mean and standard error of the trait value distribution (volume or density), $M_{true,i}$ is the “true” value of the trait estimated from the observed mean values and their standard error, $\alpha$ is the intercept, $\beta$ is the effect size (slope), and $\sigma$ is the standard deviation of the slope. Each model included four chains run for 4000 iterations, with 2000 warm-up and 2000 sampling iterations, resulting in a distribution of 8000 draws for the effect size of each trait on species richness.

*Placement of missing taxa into the Sulawesi murine phylogeny*- Four species used in this analysis do not have DNA sequence data available and were placed in the murine time-calibrated phylogeny of Nations et al. 2021 based on previous morphological analyses. *Maxomys wattsi* is thought to be closely aligned to *Maxomys rajah*, a species found across mainland Southeast Asia and the Sunda shelf (Achmadi 2010). As *Maxomys rajah* is not found on Sulawesi, we changed the name of *M. rajah* to *M. wattsi* prior to pruning the non-Sulawesi taxa from the tree. Previous morphological studies concluded that *Rattus bontanus* is in the *Rattus xanthurus* species group, *Rattus mollicomulus* is sister to *Rattus hoffmanni*, and *Rattus marmosurus* is very closely related to *Rattus xanthurus* and was previously reported as a subspecies (Musser and Carlton 2005). Using the bind.tip() function in phytools (Revell 2012), we added *Rattus bontanus* onto the *Rattus xanthurus* branch at 1mya, and *Rattus marmosurus* on the *Rattus xanthurus* branch at 0.2mya, and *Rattus bontanus* onto the *Rattus hoffmanni* branch at 1mya, resulting in a tree with 39 tips (Figure S1).

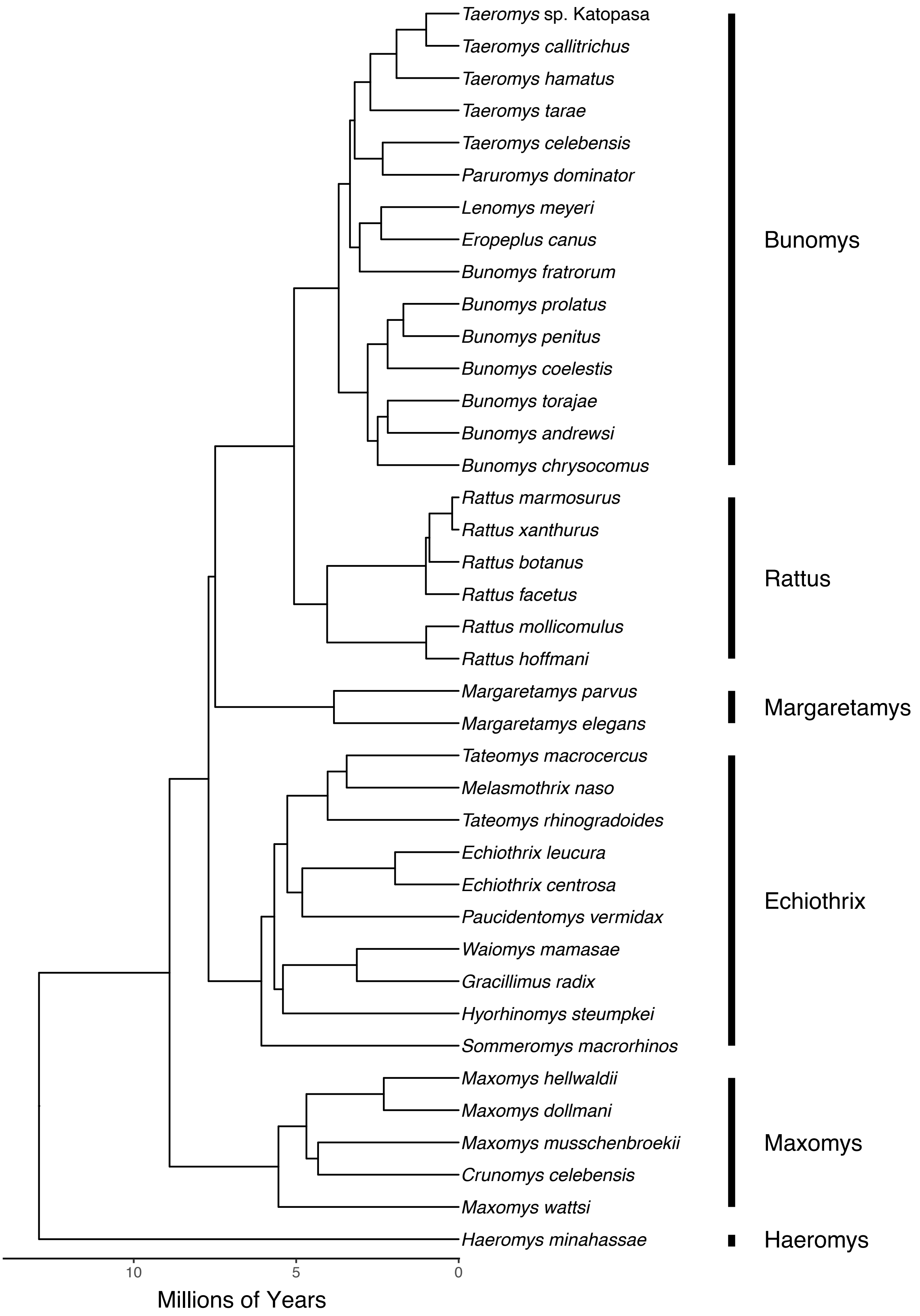


**Figure S1:** Phylogenetic hypothesis of Sulawesi murines used to estimate Phylogenetic Diversity. X-axis represents millions of years. Each of the 6 clades are labeled with black bars.

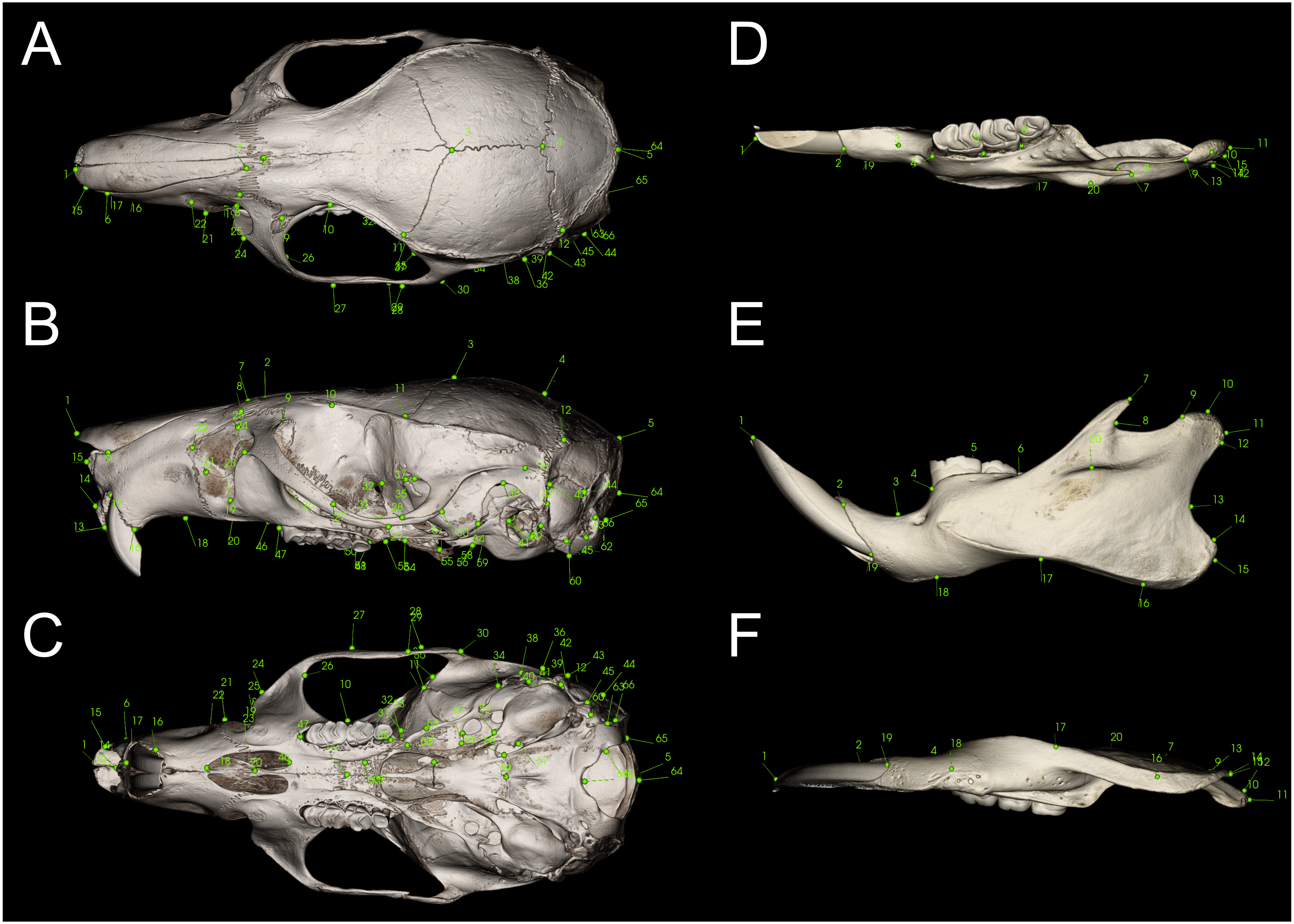


**Figure S2**: 67 Cranial (A-C) and 20 Dentary (D-F) landmarks placed on the skulls of the specimens in this study. The 67 cranial landmarks are shown in the dorsal (A), lateral (B) and ventral (C) views, and the 20 dentary landmarks are shown in the dorsal (D), lateral (E), and ventral (F) views. Image shows landmarks on LSUMZ 39095, *Maxomys musschenbroekii*.

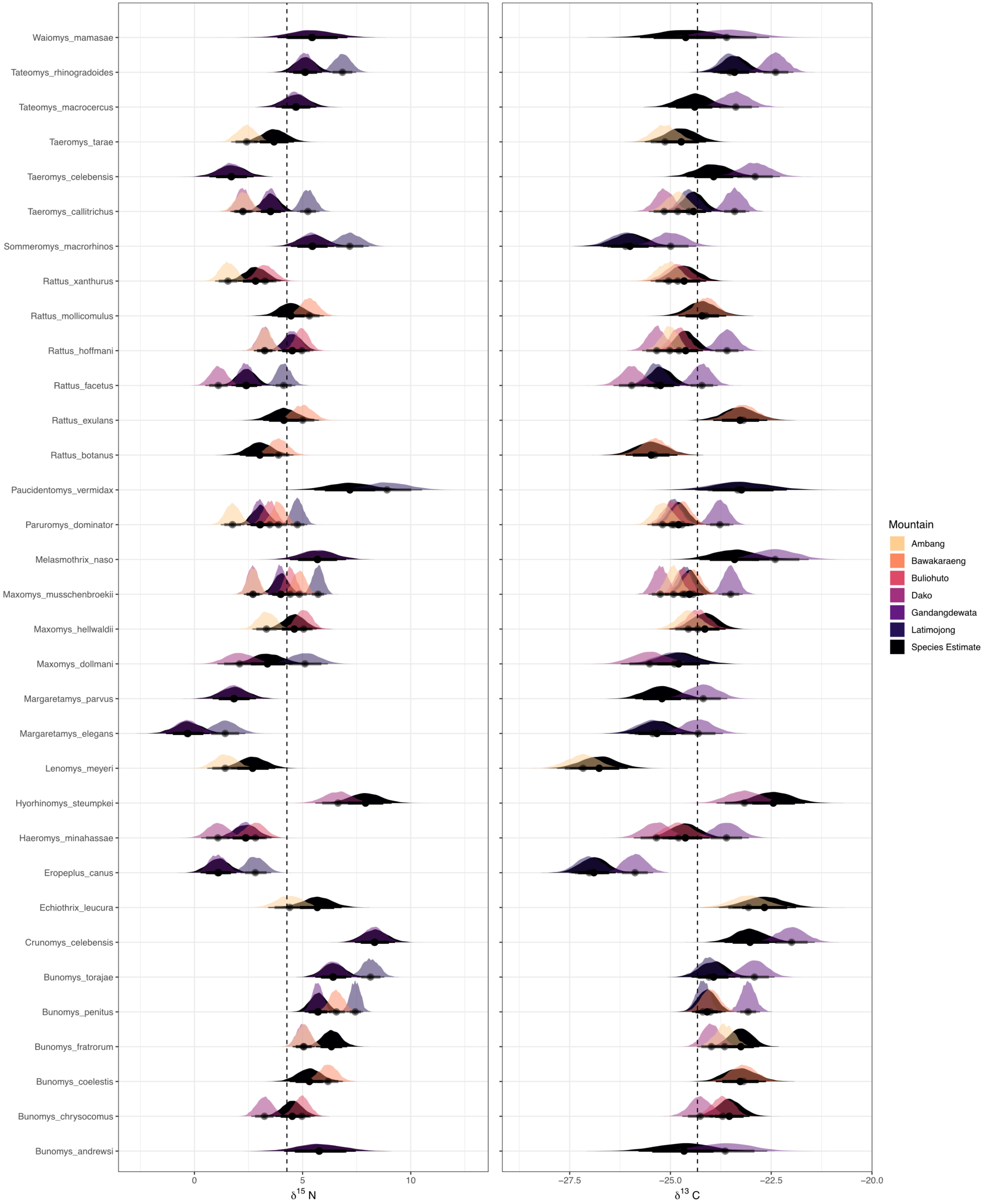


**Figure S3**: Results from multilevel model of stable isotope values that includes 'community' as a group-level effect. Using partial-pooling, a global mean for each species is estimated from the predictions of the isotope values from each species in each community. Communities are in color, with 50% alpha values, and estimated species mean values, conditioned on community, are dark with 100% alpha. The species' means are shifted by the model towards the global means shown in the vertical dashed lines. If a community often has a higher value than other communities, the model shifts the species' global mean down. For example, Bawakaraeng has high $\delta^{15}$N values, so the global means of both *Rattus mollicomulus* and *Rattus botanus* are shifted down.

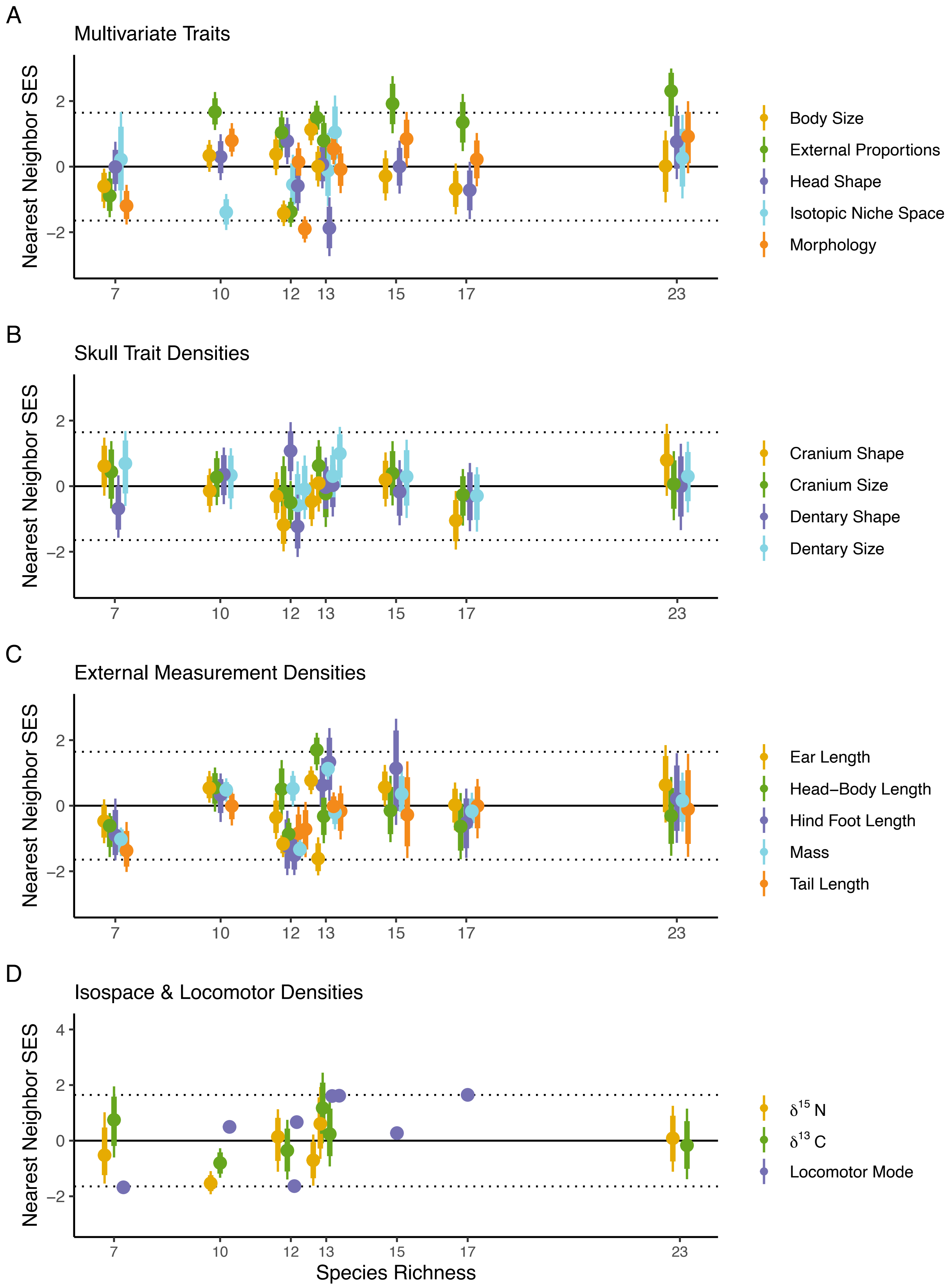

A
Multivariate Traits
Nearest Neighbor SES
2
0
−2
7
10
12
13
15
17
23
Body Size
External Proportions
Head Shape
Isotopic Niche Space
Morphology
B
Skull Trait Densities
Nearest Neighbor SES
Cranium Shape
Cranium Size
Dentary Shape
Dentary Size
C
External Measurement Densities
Nearest Neighbor SES
Ear Length
Head−Body Length
Hind Foot Length
Mass
Tail Length
D
Isospace & Locomotor Densities
Nearest Neighbor SES
4
δ15 N
δ13 C
Locomotor Mode
Species Richness

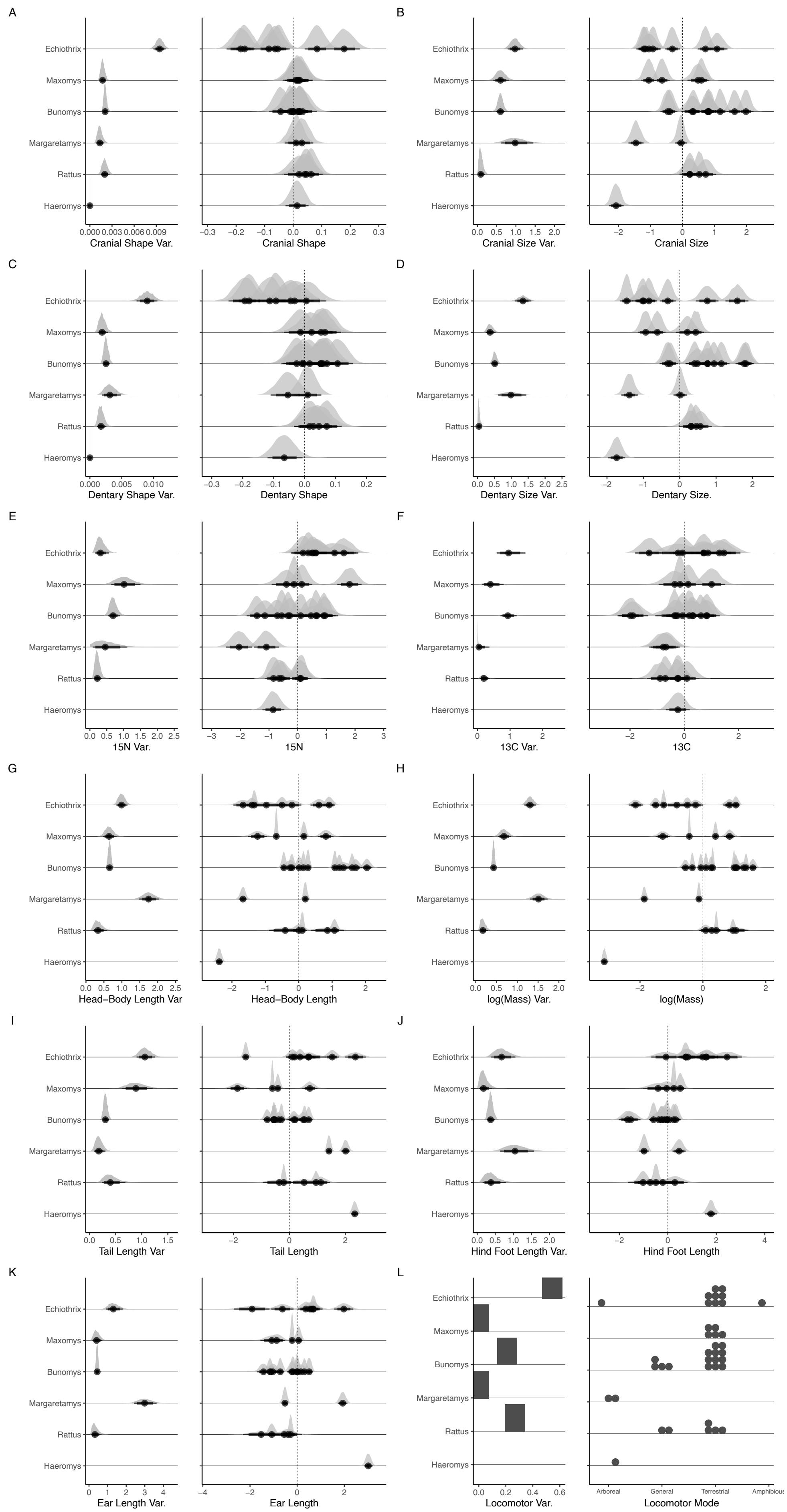
A
B
C
D
E
F
G
H
I
J
K
L
Echiothrix
Maxomys
Bunomys
Margaretamys
Rattus
Haeromys
Cranial Shape Var.
Cranial Shape
Cranial Size Var.
Cranial Size
Dentary Shape Var.
Dentary Shape
Dentary Size Var.
Dentary Size.
15N Var.
15N
13C Var.
13C
Head−Body Length Var
Head−Body Length
log(Mass) Var.
log(Mass)
Tail Length Var
Tail Length
Hind Foot Length Var.
Hind Foot Length
Ear Length Var.
Ear Length
Locomotor Var.
Locomotor Mode
Arboreal
General
Terrestrial
Amphibious

**Supporting Table 1**: Description of the 12 traits and five combined traits and the methods used to calculate functional space and density.

| Individual Traits | Data Description | Functional Space Calculation | No. communities sampled |
|---|---|---|---|
| Cranium Shape | 67 3D geometric morphometric landmarks on 64 $\mu$CT scans of crania from 38 species (n=1-3, mean = 1.9). | The posterior distribution of predicted Generalized Procrustes Analysis values was estimated for each axis for each species. Species were grouped by community, then, for each draw from the posterior, the sum of variance was calculated for each community from 36 axes (>95% of PC variation). A nearest neighbor value was calculated for each draw from the 36 PC axes. | 9 |
| Cranium Size | Unit-centroid size calculated from 64 $\mu$CT scans of crania from 38 species (n=1-3, mean = 1.9). | We estimated the posterior distribution of the predicted values for each species, sorted the species predictions by community, then estimated the variance and nearest neighbor density for each posterior draw. | 9 |
| Dentary Shape | 20 3D geometric morphometric landmarks on 61 samples from 36 species (n=1-3, mean = 1.7). | Species estimates and community sum of variances were estimated as in Cranial Shape above using all 20 axes (>95% of PC variation). A nearest neighbor value was calculated for each draw from the 20 PC axes for each community. | 9 |
| Dentary Size | Unit-centroid size calculated from 61 $\mu$CT scans of dentaries from 36 species (n=1-3, mean = 1.7). | Processed as Cranium Size above | 9 |
| Head-Body Length | Head-Body length (in mm) for 630 specimens of 39 species (n=1-95, mean = 16). | Processed as Cranium Size above | 9 |
| Relative Tail Length | We calculated Tail Length / Head-Body Length for 630 specimens of 39 species (n=1-95, mean = 16). | Processed as Cranium Size above | 9 |
| Relative Ear Length | Ear Length / Head-Body Length for 630 specimens of 39 species (n=1-95, mean = 16). | Processed as Cranium Size above | 9 |

| | | | |
|---|---|---|---|
| Relative Hind Foot Length | Hind Foot Length / Head-Body Length for 630 specimens of 39 species (n=1-95, mean = 16). | Processed as Cranium Size above | 9 |
| Log(Mass) | Natural log of Mass in grams for 630 specimens of 39 species (n=1-95, mean = 16). | Processed as Cranium Size above | 9 |
| $\delta^{13}C$ | $\delta^{13}C$ values from hair samples of 286 specimens of 34 species (n=1-24, mean = 8.7). | Species' estimates were processed as Cranium Shape above. For randomization analyses, species' predicted values were estimated using a second multilevel model with mountain community as an additional group-level effect, resulting in a posterior distribution for each species lacking any geographic influence. (Figure S3). | 6 |
| $\delta^{15}N$ | $\delta^{15}N$ values from hair samples of 286 specimens of 34 species (n=1-24, mean = 8.7). | Processed as $\delta^{13}C$. | 6 |
| Locomotor Mode | 39 species were assigned 1 of 4 locomotor modes: Arboreal, General, Terrestrial, and Amphibious. | Ordinal locomotor variables were coded 1-4, grouped species by community, then estimated the variance of each community. Locomotor scores were transformed into a binary array, a redundancy analysis (rda) was performed, then the nearest neighbor scores were estimated from the rda PC scores. | 9 |

**Combined Traits**

| | | | |
|---|---|---|---|
| Morphology | Posterior predictions of Cranial Shape, Cranial Size, Dentary Shape, Dentary Size, Head-Body Length, Tail Length, Ear Length, Hind Foot Length, and Mass combined into a matrix. | Posterior predictions of each trait were grouped by community, then the sum of variance and nearest neighbor density were calculated in this 9-dimensional space for each posterior draw. | 9 |
| External Proportions | Posterior predictions of Head-Body Length, Tail Length, Ear Length, Hind Foot Length, and Mass combined into a matrix. | 5-dimensional space, processed as Morphology above. | 9 |

| | | | |
|---|---|---|---|
| Body Size | Posterior predictions of Cranial Size, Dentary Size, Head-Body Length, and Mass combined into a matrix. | 4-dimensional space, processed as Morphology above. | 9 |
| Head Shape | Posterior predictions of 36 Cranial PC axes and the 20 Dentary PC axes combined into one matrix of 56 columns. | We grouped posterior predictions by community, then calculated the sum of variance for each posterior draw with each of the 56 axes. We estimated the nearest neighbor density from the 56-dimensional matrix. | 9 |
| Isotopic Niche Space | Posterior predictions of $\delta^{15}N$ and $\delta^{13}C$ values, conditioned on locality, were combined into a 2-dimensional matrix. | 2-dimensional space, processed as Morphology above. | 6 |

**Supporting Table 2**: Mean values of total ecological volume along with each of the five trait axes, with lower and upper 89% probability intervals in parentheses. Locomotor mode variance was directly estimated using the number of discrete locomotor modes (Arboreal, General, Terrestrial, and Amphibious) in each community.

| Trait | Community | Volume | Volume SES | Density | Density SES |
|---|---|---|---|---|---|
| Body Size | Ambang | 1.4 (1.34, 1.45) | 0.16 (0, 0.33) | 0.5 (0.46, 0.54) | 1.21 (0.77, 1.64) |
| | Bawakaraeng | 0.53 (0.45, 0.63) | -1.19 (-1.32, -1.03) | 0.48 (0.34, 0.6) | -0.51 (-1.12, 0.01) |
| | Buliohuto | 1.56 (1.5, 1.63) | 0.51 (0.37, 0.66) | 0.53 (0.47, 0.59) | 0.27 (-0.19, 0.73) |
| | Dako | 1.62 (1.56, 1.69) | 0.88 (0.7, 1.06) | 0.46 (0.4, 0.51) | 0.43 (-0.2, 1.07) |
| | Gandangdewata | 1.4 (1.34, 1.46) | 1.44 (0.96, 1.92) | 0.27 (0.23, 0.3) | -0.06 (-1.07, 0.96) |
| | Katopasa | 0.39 (0.36, 0.41) | -2.56 (-2.63, -2.49) | 0.25 (0.21, 0.29) | -2.04 (-2.49, -1.56) |
| | Latimojong | 0.93 (0.85, 1.01) | -1.08 (-1.31, -0.84) | 0.4 (0.35, 0.45) | 0.04 (-0.53, 0.62) |
| | Nokilalaki | 1.35 (1.3, 1.41) | 0.48 (0.29, 0.68) | 0.3 (0.26, 0.34) | -0.74 (-1.52, 0.07) |
| | Torompupu | 1.45 (1.38, 1.52) | 0.54 (0.31, 0.77) | 0.34 (0.29, 0.39) | -0.24 (-0.96, 0.55) |
| External Proportions | Ambang | 1.13 (1.07, 1.18) | 0.01 (-0.18, 0.21) | 1.3 (1.22, 1.37) | 1.5 (0.98, 1.98) |
| | Bawakaraeng | 0.42 (0.33, 0.52) | -1.16 (-1.33, -0.94) | 0.95 (0.74, 1.18) | -1.13 (-1.83, -0.41) |
| | Buliohuto | 1.21 (1.16, 1.27) | 0.39 (0.25, 0.54) | 1.49 (1.39, 1.59) | 1.75 (1.23, 2.29) |
| | Dako | 1.31 (1.25, 1.38) | 0.76 (0.55, 0.99) | 1.27 (1.18, 1.37) | 1.14 (0.58, 1.7) |
| | Gandangdewata | 1.25 (1.19, 1.32) | 1.58 (1.01, 2.19) | 1.16 (1.06, 1.26) | 2.44 (1.3, 3.55) |
| | Katopasa | 0.35 (0.32, 0.38) | -2.47 (-2.57, -2.36) | 0.82 (0.74, 0.9) | -1.89 (-2.36, -1.41) |
| | Latimojong | 0.77 (0.7, 0.85) | -1.16 (-1.43, -0.87) | 1.17 (1.07, 1.28) | 0.68 (-0.01, 1.38) |
| | Nokilalaki | 1.17 (1.12, 1.23) | 0.56 (0.33, 0.8) | 1.17 (1.07, 1.26) | 1.08 (0.34, 1.79) |
| | Torompupu | 1.31 (1.24, 1.39) | 0.84 (0.56, 1.14) | 1.3 (1.19, 1.4) | 2.18 (1.31, 3.06) |
| Isospace | Ambang | 0.52 (0.4, 0.66) | -0.22 (-0.95, 0.57) | 0.46 (0.34, 0.57) | -0.05 (-1.25, 1.15) |
| | Bawakaraeng | 0.34 (0.22, 0.49) | -0.12 (-0.74, 0.63) | 0.54 (0.38, 0.72) | 0.6 (-0.83, 2.11) |
| | Buliohuto | 0.18 (0.12, 0.24) | -1.67 (-1.97, -1.33) | 0.39 (0.29, 0.48) | -0.66 (-1.35, 0.04) |
| | Dako | 0.55 (0.42, 0.7) | 0.03 (-0.75, 0.88) | 0.38 (0.28, 0.49) | -0.69 (-1.67, 0.34) |
| | Gandangdewata | 0.77 (0.65, 0.9) | 0.5 (-0.37, 1.46) | 0.4 (0.32, 0.47) | -0.06 (-1.37, 1.28) |
| | Latimojong | 0.81 (0.64, 0.99) | 1.11 (0.27, 2.07) | 0.53 (0.41, 0.65) | 0.69 (-0.45, 1.86) |
| Morphology | Ambang | 0.93 (0.89, 0.98) | -0.07 (-0.27, 0.13) | 2.08 (1.96, 2.21) | 0.63 (0.25, 1.03) |
| | Bawakaraeng | 0.38 (0.31, 0.45) | -1.18 (-1.35, -0.99) | 1.04 (0.84, 1.27) | -2.25 (-2.69, -1.77) |
| | Buliohuto | 1 (0.96, 1.05) | 0.37 (0.22, 0.52) | 2.33 (2.19, 2.48) | 0.78 (0.39, 1.19) |
| | Dako | 1.11 (1.06, 1.16) | 0.88 (0.65, 1.11) | 2.36 (2.22, 2.5) | 1.3 (0.83, 1.77) |
| | Gandangdewata | 1.08 (1.04, 1.13) | 1.78 (1.2, 2.38) | 1.78 (1.66, 1.91) | 0.96 (0.21, 1.71) |
| | Katopasa | 0.32 (0.29, 0.34) | -2.47 (-2.57, -2.36) | 1.24 (1.1, 1.38) | -2.24 (-2.68, -1.78) |
| | Latimojong | 0.72 (0.67, 0.78) | -1.15 (-1.4, -0.88) | 1.64 (1.49, 1.8) | -0.7 (-1.2, -0.16) |
| | Nokilalaki | 1.01 (0.97, 1.05) | 0.59 (0.37, 0.82) | 1.95 (1.82, 2.09) | 0.57 (0, 1.17) |
| | Torompupu | 1.08 (1.02, 1.13) | 0.74 (0.47, 1.03) | 2.04 (1.9, 2.19) | 0.9 (0.38, 1.44) |
| Carbon13 | Ambang | 0.64 (0.42, 0.88) | 0.73 (-0.32, 1.9) | 0.22 (0.14, 0.29) | 0.86 (-0.44, 2.17) |
| | Bawakaraeng | 0.35 (0.18, 0.58) | 0.18 (-0.54, 1.11) | 0.3 (0.18, 0.4) | 1.43 (-0.16, 2.9) |
| | Buliohuto | 0.17 (0.09, 0.26) | -1.13 (-1.48, -0.71) | 0.17 (0.12, 0.22) | -0.39 (-0.94, 0.16) |
| | Dako | 0.48 (0.3, 0.69) | 0.09 (-0.75, 1.08) | 0.11 (0.06, 0.18) | -0.78 (-1.56, 0.15) |
| | Gandangdewata | 0.58 (0.42, 0.76) | -0.3 (-1.22, 0.76) | 0.1 (0.06, 0.14) | -0.43 (-1.69, 0.95) |
| | Latimojong | 0.68 (0.46, 0.94) | 0.73 (-0.21, 1.89) | 0.18 (0.11, 0.26) | 0.22 (-0.93, 1.53) |
| Cranium Shape | Ambang | 0.39 (0.35, 0.43) | -0.72 (-1.02, -0.4) | 0.49 (0.44, 0.54) | -0.37 (-0.79, 0.06) |
| | Bawakaraeng | 0.32 (0.25, 0.39) | -0.51 (-0.86, -0.14) | 0.63 (0.49, 0.77) | -0.46 (-0.99, 0.14) |
| | Buliohuto | 0.42 (0.37, 0.46) | -0.25 (-0.53, 0.03) | 0.67 (0.6, 0.74) | 0.38 (-0.04, 0.79) |
| | Dako | 0.42 (0.38, 0.47) | -0.45 (-0.82, -0.08) | 0.58 (0.51, 0.65) | 0.06 (-0.4, 0.51) |
| | Gandangdewata | 0.7 (0.65, 0.75) | 1.61 (1.07, 2.18) | 0.46 (0.41, 0.51) | -0.04 (-0.78, 0.68) |
| | Katopasa | 0.22 (0.19, 0.26) | -1.82 (-2.07, -1.54) | 0.42 (0.33, 0.5) | -1.42 (-2.03, -0.82) |
| | Latimojong | 0.67 (0.62, 0.72) | 1.36 (0.97, 1.77) | 0.68 (0.59, 0.76) | 0.93 (0.37, 1.44) |
| | Nokilalaki | 0.49 (0.45, 0.52) | -0.24 (-0.57, 0.09) | 0.39 (0.34, 0.45) | -0.89 (-1.44, -0.35) |
| | Torompupu | 0.47 (0.43, 0.51) | -0.28 (-0.62, 0.06) | 0.5 (0.44, 0.56) | -0.2 (-0.72, 0.31) |

| Trait | Community | Volume | Volume SES | Density | Density SES |
|---|---|---|---|---|---|
| Cranium Size | Ambang | 1.05 (0.91, 1.19) | -0.27 (-0.91, 0.4) | 0.28 (0.23, 0.32) | 0.87 (0.02, 1.63) |
| | Bawakaraeng | 1.08 (0.84, 1.31) | -0.4 (-0.99, 0.2) | 0.5 (0.38, 0.62) | 0.09 (-0.89, 1.02) |
| | Buliohuto | 1.26 (1.08, 1.44) | 0.38 (-0.21, 0.98) | 0.35 (0.29, 0.41) | 0.13 (-0.56, 0.84) |
| | Dako | 1.34 (1.17, 1.52) | 0.85 (0.19, 1.53) | 0.26 (0.19, 0.32) | -0.4 (-1.46, 0.61) |
| | Gandangdewata | 1.01 (0.9, 1.13) | 0.33 (-0.66, 1.34) | 0.14 (0.12, 0.16) | -0.32 (-1.29, 0.63) |
| | Katopasa | 0.67 (0.54, 0.8) | -1.86 (-2.36, -1.34) | 0.24 (0.19, 0.29) | -0.84 (-1.63, 0.02) |
| | Latimojong | 0.67 (0.56, 0.79) | -1.75 (-2.24, -1.24) | 0.23 (0.19, 0.27) | -0.42 (-1.12, 0.22) |
| | Nokilalaki | 1.04 (0.91, 1.17) | 0.12 (-0.5, 0.74) | 0.19 (0.16, 0.22) | 0.07 (-0.91, 0.96) |
| | Torompupu | 1.11 (0.97, 1.24) | 0.31 (-0.33, 0.97) | 0.23 (0.19, 0.27) | -0.06 (-0.91, 0.81) |
| Dentary Shape | Ambang | 0.47 (0.38, 0.57) | -0.24 (-0.83, 0.34) | 0.8 (0.68, 0.92) | 0.11 (-0.57, 0.78) |
| | Bawakaraeng | 0.21 (0.14, 0.3) | -0.92 (-1.19, -0.61) | 0.67 (0.43, 0.92) | -0.57 (-1.44, 0.39) |
| | Buliohuto | 0.52 (0.41, 0.64) | 0.25 (-0.31, 0.84) | 0.93 (0.78, 1.06) | 0.33 (-0.32, 0.95) |
| | Dako | 0.77 (0.64, 0.89) | 1.35 (0.72, 2.01) | 1.02 (0.89, 1.15) | 1.16 (0.44, 1.86) |
| | Gandangdewata | 0.58 (0.51, 0.65) | 0.01 (-0.52, 0.55) | 0.71 (0.61, 0.81) | -0.13 (-1.1, 0.83) |
| | Katopasa | 0.5 (0.43, 0.56) | 0.14 (-0.23, 0.53) | 0.52 (0.38, 0.67) | -1.45 (-2.18, -0.71) |
| | Latimojong | 0.23 (0.17, 0.29) | -1.67 (-2.01, -1.29) | 0.77 (0.65, 0.9) | -0.16 (-0.77, 0.47) |
| | Nokilalaki | 0.65 (0.57, 0.73) | 0.69 (0.19, 1.19) | 0.76 (0.64, 0.88) | -0.14 (-0.99, 0.73) |
| | Torompupu | 0.47 (0.39, 0.55) | -0.27 (-0.8, 0.28) | 0.77 (0.66, 0.88) | 0.02 (-0.63, 0.65) |
| Dentary Size | Ambang | 0.92 (0.81, 1.04) | -0.13 (-0.64, 0.41) | 0.48 (0.44, 0.51) | 1.67 (1.43, 1.89) |
| | Bawakaraeng | 0.86 (0.68, 1.06) | -0.59 (-1.12, -0.04) | 0.35 (0.25, 0.43) | 0.03 (-0.52, 0.48) |
| | Buliohuto | 0.95 (0.82, 1.09) | -0.08 (-0.53, 0.4) | 0.22 (0.18, 0.26) | -0.13 (-0.51, 0.21) |
| | Dako | 1.17 (1.03, 1.31) | 0.8 (0.26, 1.36) | 0.15 (0.09, 0.2) | -0.49 (-1.03, 0.02) |
| | Gandangdewata | 0.95 (0.85, 1.05) | 0.36 (-0.36, 1.1) | 0.09 (0.07, 0.1) | -1.11 (-1.28, -0.94) |
| | Katopasa | 0.52 (0.42, 0.62) | -1.92 (-2.35, -1.45) | 0.16 (0.13, 0.2) | -0.48 (-0.72, -0.22) |
| | Latimojong | 0.73 (0.63, 0.84) | -0.85 (-1.28, -0.4) | 0.19 (0.16, 0.22) | -0.17 (-0.41, 0.05) |
| | Nokilalaki | 0.94 (0.82, 1.05) | 0.03 (-0.54, 0.62) | 0.36 (0.33, 0.38) | 1.26 (1.04, 1.46) |
| | Torompupu | 0.97 (0.86, 1.09) | 0.49 (-0.05, 1.07) | 0.14 (0.11, 0.17) | -0.61 (-0.84, -0.4) |
| Ear Length | Ambang | 1.39 (1.23, 1.56) | 0.57 (0.21, 0.96) | 0.3 (0.27, 0.34) | 0.85 (0.42, 1.3) |
| | Bawakaraeng | 0.43 (0.17, 0.78) | -0.62 (-0.97, -0.15) | 0.25 (0.12, 0.37) | -0.64 (-1.32, 0.04) |
| | Buliohuto | 1.45 (1.28, 1.63) | 0.72 (0.41, 1.05) | 0.37 (0.32, 0.43) | 0.76 (0.34, 1.22) |
| | Dako | 1.73 (1.51, 1.97) | 1.34 (0.85, 1.84) | 0.24 (0.18, 0.29) | -0.26 (-0.89, 0.44) |
| | Gandangdewata | 1.25 (1.09, 1.44) | 0.96 (0.08, 1.93) | 0.17 (0.14, 0.21) | 0.87 (-0.41, 2.25) |
| | Katopasa | 0.26 (0.2, 0.34) | -1.86 (-2, -1.71) | 0.12 (0.08, 0.16) | -1.57 (-2.03, -1.16) |
| | Latimojong | 0.32 (0.21, 0.45) | -1.89 (-2.17, -1.56) | 0.11 (0.06, 0.15) | -1.56 (-2.12, -0.95) |
| | Nokilalaki | 1.22 (1.08, 1.37) | 0.45 (0.04, 0.89) | 0.2 (0.17, 0.24) | -0.18 (-0.96, 0.57) |
| | Torompupu | 1.09 (0.95, 1.24) | 0.02 (-0.35, 0.41) | 0.25 (0.22, 0.29) | 0.57 (-0.05, 1.26) |
| Head Shape | Ambang | 0.49 (0.43, 0.55) | -0.43 (-0.8, -0.06) | 1.2 (1.08, 1.32) | 0.18 (-0.39, 0.72) |
| | Bawakaraeng | 0.32 (0.26, 0.38) | -0.85 (-1.11, -0.56) | 1.1 (0.82, 1.4) | -0.43 (-1.24, 0.45) |
| | Buliohuto | 0.52 (0.46, 0.59) | 0.11 (-0.28, 0.5) | 1.35 (1.21, 1.5) | 0.33 (-0.24, 0.88) |
| | Dako | 0.67 (0.6, 0.73) | 0.66 (0.2, 1.13) | 1.44 (1.31, 1.58) | 1.05 (0.43, 1.7) |
| | Gandangdewata | 0.69 (0.64, 0.74) | 0.88 (0.39, 1.4) | 1.1 (0.99, 1.21) | 0.14 (-0.8, 1.12) |
| | Katopasa | 0.27 (0.23, 0.32) | -1.01 (-1.3, -0.7) | 0.82 (0.67, 0.99) | -1.76 (-2.47, -1.01) |
| | Latimojong | 0.63 (0.59, 0.68) | -0.41 (-0.68, -0.11) | 1.14 (1, 1.27) | -0.22 (-0.79, 0.35) |
| | Nokilalaki | 0.67 (0.61, 0.72) | 0.31 (-0.08, 0.72) | 1.07 (0.95, 1.19) | -0.51 (-1.34, 0.32) |
| | Torompupu | 0.54 (0.48, 0.59) | -0.37 (-0.81, 0.08) | 1.14 (1.02, 1.27) | -0.05 (-0.69, 0.59) |
| Head-Body Length | Ambang | 1.31 (1.23, 1.39) | 0.15 (-0.14, 0.46) | 0.3 (0.27, 0.33) | 2.04 (1.38, 2.63) |
| | Bawakaraeng | 0.65 (0.51, 0.82) | -1.23 (-1.52, -0.86) | 0.29 (0.17, 0.35) | -0.82 (-1.66, -0.35) |
| | Buliohuto | 1.38 (1.28, 1.49) | 0.32 (0.05, 0.6) | 0.31 (0.27, 0.36) | 0.69 (-0.01, 1.34) |
| | Dako | 1.44 (1.34, 1.54) | 0.71 (0.38, 1.04) | 0.26 (0.21, 0.3) | 0.51 (-0.29, 1.27) |
| | Gandangdewata | 1.34 (1.24, 1.43) | 1.49 (0.75, 2.24) | 0.11 (0.09, 0.13) | -0.47 (-1.68, 0.66) |
| | Katopasa | 0.47 (0.43, 0.52) | -2.51 (-2.65, -2.36) | 0.16 (0.13, 0.19) | -1.27 (-1.87, -0.71) |
| | Latimojong | 0.94 (0.82, 1.06) | -0.96 (-1.38, -0.53) | 0.19 (0.15, 0.23) | -0.28 (-1.09, 0.52) |
| | Nokilalaki | 1.3 (1.21, 1.39) | 0.56 (0.22, 0.89) | 0.15 (0.11, 0.18) | -0.81 (-1.84, 0.29) |
| | Torompupu | 1.34 (1.23, 1.45) | 0.5 (0.09, 0.92) | 0.18 (0.14, 0.22) | -0.16 (-1.2, 0.89) |

| Trait | Community | Volume | Volume SES | Density | Density SES |
|---|---|---|---|---|---|
| Hind Foot Length | Ambang | 0.81 (0.65, 0.99) | 0.17 (-0.41, 0.8) | 0.22 (0.17, 0.27) | 0.57 (-0.24, 1.33) |
| | Bawakaraeng | 0.24 (0.06, 0.55) | -0.94 (-1.39, -0.17) | 0.16 (0.06, 0.29) | -0.9 (-1.74, 0.16) |
| | Buliohuto | 0.63 (0.5, 0.78) | -0.09 (-0.5, 0.34) | 0.25 (0.19, 0.3) | 0.18 (-0.56, 0.84) |
| | Dako | 0.58 (0.44, 0.74) | -0.37 (-0.88, 0.23) | 0.12 (0.06, 0.19) | -1.12 (-2, -0.14) |
| | Gandangdewata | 0.96 (0.76, 1.18) | 0.76 (-0.38, 2.02) | 0.12 (0.08, 0.16) | 0.33 (-1.01, 1.69) |
| | Katopasa | 0.19 (0.11, 0.29) | -1.8 (-2.06, -1.48) | 0.09 (0.05, 0.14) | -1.79 (-2.47, -0.96) |
| | Latimojong | 0.95 (0.68, 1.26) | 0.84 (-0.17, 1.98) | 0.27 (0.2, 0.34) | 1.5 (0.37, 2.58) |
| | Nokilalaki | 0.79 (0.63, 0.96) | 0.24 (-0.41, 0.92) | 0.13 (0.09, 0.18) | -0.5 (-1.5, 0.53) |
| | Torompupu | 0.96 (0.73, 1.23) | 0.72 (-0.12, 1.68) | 0.23 (0.14, 0.31) | 1.29 (-0.52, 2.94) |
| Mass | Ambang | 1.49 (1.42, 1.56) | 0.16 (-0.02, 0.34) | 0.32 (0.3, 0.34) | 1.32 (0.98, 1.66) |
| | Bawakaraeng | 0.42 (0.33, 0.52) | -1.13 (-1.24, -0.99) | 0.21 (0.12, 0.27) | -1.23 (-1.71, -0.84) |
| | Buliohuto | 1.75 (1.66, 1.84) | 0.62 (0.47, 0.78) | 0.36 (0.33, 0.4) | 0.69 (0.3, 1.06) |
| | Dako | 1.81 (1.72, 1.9) | 0.96 (0.76, 1.16) | 0.3 (0.26, 0.34) | 0.54 (0.05, 1.01) |
| | Gandangdewata | 1.46 (1.38, 1.53) | 1.19 (0.66, 1.72) | 0.14 (0.12, 0.16) | -0.1 (-1, 0.78) |
| | Katopasa | 0.3 (0.27, 0.32) | -2.49 (-2.55, -2.43) | 0.12 (0.1, 0.15) | -1.95 (-2.34, -1.57) |
| | Latimojong | 0.92 (0.81, 1.03) | -1.1 (-1.35, -0.84) | 0.22 (0.19, 0.25) | -0.13 (-0.58, 0.29) |
| | Nokilalaki | 1.41 (1.34, 1.48) | 0.4 (0.2, 0.61) | 0.19 (0.16, 0.21) | -0.32 (-0.88, 0.28) |
| | Torompupu | 1.56 (1.47, 1.66) | 0.54 (0.29, 0.8) | 0.23 (0.19, 0.26) | 0.34 (-0.41, 1.07) |
| Nitrogen15 | Ambang | 0.4 (0.29, 0.54) | -0.98 (-1.49, -0.4) | 0.14 (0.08, 0.19) | -0.69 (-1.58, 0.25) |
| | Bawakaraeng | 0.33 (0.19, 0.54) | -0.36 (-0.93, 0.47) | 0.2 (0.1, 0.31) | -0.13 (-1.38, 1.35) |
| | Buliohuto | 0.19 (0.12, 0.27) | -1.5 (-1.79, -1.16) | 0.13 (0.08, 0.19) | -1.13 (-1.66, -0.56) |
| | Dako | 0.63 (0.45, 0.83) | -0.05 (-0.81, 0.82) | 0.16 (0.09, 0.24) | -0.41 (-1.63, 0.82) |
| | Gandangdewata | 0.96 (0.79, 1.16) | 1.03 (0.07, 2.13) | 0.11 (0.08, 0.15) | -0.35 (-1.57, 0.93) |
| | Latimojong | 0.93 (0.71, 1.2) | 1.05 (0.13, 2.13) | 0.18 (0.11, 0.25) | -0.03 (-1.18, 1.21) |
| Tail Length | Ambang | 0.64 (0.58, 0.7) | -1.43 (-1.65, -1.21) | 0.19 (0.16, 0.21) | 0 (-0.45, 0.43) |
| | Bawakaraeng | 0.35 (0.21, 0.54) | -1.19 (-1.5, -0.75) | 0.14 (0.06, 0.24) | -1.53 (-2.22, -0.74) |
| | Buliohuto | 0.86 (0.77, 0.96) | -0.26 (-0.51, -0.01) | 0.24 (0.2, 0.28) | 0.09 (-0.48, 0.62) |
| | Dako | 1 (0.91, 1.09) | -0.13 (-0.43, 0.16) | 0.16 (0.1, 0.2) | -0.75 (-1.64, 0.09) |
| | Gandangdewata | 1.26 (1.15, 1.38) | 1.12 (0.42, 1.85) | 0.11 (0.08, 0.15) | -0.12 (-1.6, 1.51) |
| | Katopasa | 0.51 (0.4, 0.62) | -1.63 (-1.96, -1.29) | 0.15 (0.1, 0.2) | -0.93 (-1.79, -0.11) |
| | Latimojong | 0.73 (0.61, 0.87) | -1.1 (-1.53, -0.64) | 0.18 (0.13, 0.22) | -0.27 (-1.08, 0.5) |
| | Nokilalaki | 1.15 (1.05, 1.25) | 0.64 (0.25, 1.04) | 0.16 (0.13, 0.19) | 0.02 (-0.84, 0.79) |
| | Torompupu | 1.6 (1.42, 1.79) | 2.08 (1.41, 2.79) | 0.16 (0.1, 0.23) | -0.13 (-1.48, 1.34) |
| Locomotion | Ambang | 0.44 | -0.66 | 0.38 | 1.41 |
| | Bawakaraeng | 0.14 | -1.49 | 0.2 | -1.2 |
| | Buliohuto | 0.49 | -0.38 | 0.38 | 1.31 |
| | Dako | 0.64 | 0.41 | 0.38 | 1.31 |
| | Gandangdewata | 0.72 | 1.96 | 0.45 | 3.98 |
| | Katopasa | 0.22 | -1.84 | 0.25 | -0.61 |
| | Latimojong | 0.6 | 0.4 | 0.38 | 1.48 |
| | Nokilalaki | 0.63 | 0.55 | 0.37 | 1.67 |
| | Torompupu | 0.58 | 0.35 | 0.33 | 0.79 |
| Phylogenetic Diversity | Ambang | 71.078 | -0.18 | | |
| | Bawakaraeng | 35.59 | -1.34 | | |
| | Buliohuto | 64.133 | 0.57 | | |
| | Dako | 76.681 | 1.09 | | |
| | Gandangdewata | 118.096 | 1.26 | | |
| | Katopasa | 53.371 | -2.05 | | |
| | Latimojong | 71.176 | -0.24 | | |
| | Nokilalaki | 88.204 | 0.49 | | |
| | Torompupu | 87.708 | 0.98 | | |

**Table S3**: Results from regression analyses. Beta is the slope of the linear regression. $R^2$ is the Bayesian $R^2$. Mean values are reported with lower and upper 89% probability intervals in parentheses. Non-zero Beta probabilities are in bold.

| **Volume** | **Beta** | **Intercept** | **sigma** | **$R^2$** |
|---|---|---|---|---|
| Body Size | 0.048 (-0.054, 0.148) | -1.121 (-2.531, 0.34) | 0.823 (0.58, 1.132) | 0.122 (0.001, 0.344) |
| Carbon 13 Isotope | 0.003 (-0.094, 0.099) | -0.005 (-1.307, 1.32) | 0.722 (0.471, 1.062) | 0.111 (0.001, 0.321) |
| Cranial Shape | **0.142 (0.024, 0.261)** | -2.014 (-3.649, -0.349) | 0.923 (0.672, 1.246) | 0.294 (0.017, 0.539) |
| Cranial Size | 0.058 (-0.049, 0.166) | -0.979 (-2.502, 0.516) | 0.883 (0.637, 1.199) | 0.123 (0.001, 0.349) |
| Dentary Shape | 0.046 (-0.065, 0.16) | -0.713 (-2.279, 0.839) | 0.915 (0.668, 1.232) | 0.1 (0.001, 0.31) |
| Dentary Size | 0.061 (-0.041, 0.162) | -1.044 (-2.455, 0.399) | 0.815 (0.582, 1.115) | 0.146 (0.002, 0.385) |
| Ear Length | 0.084 (-0.047, 0.214) | -1.198 (-3, 0.627) | 1.036 (0.764, 1.371) | 0.138 (0.001, 0.368) |
| External Proportions | 0.043 (-0.058, 0.144) | -1.455 (-2.856, 0) | 0.815 (0.569, 1.131) | 0.118 (0.001, 0.345) |
| Head Shape | 0.096 (-0.017, 0.206) | -1.423 (-2.957, 0.179) | 0.897 (0.648, 1.223) | 0.196 (0.004, 0.443) |
| Head-Body Length | **0.153 (0.034, 0.271)** | -2.212 (-3.843, -0.556) | 0.943 (0.687, 1.272) | 0.309 (0.022, 0.545) |
| Hind Foot Length | **0.108 (0.012, 0.207)** | -1.54 (-2.937, -0.16) | 0.788 (0.556, 1.093) | 0.273 (0.013, 0.523) |
| Isospace | 0.08 (-0.029, 0.186) | -1.052 (-2.505, 0.438) | 0.824 (0.563, 1.177) | 0.223 (0.003, 0.494) |
| Locomotor Mode | **0.188 (0.100, 0.269)** | -2.497 (-3.686, -1.292) | 0.672 (0.467, 0.948) | 0.586 (0.263, 0.732) |
| Mass | **0.128 (0.007, 0.249)** | -1.827 (-3.546, -0.117) | 0.98 (0.719, 1.315) | 0.234 (0.008, 0.476) |
| Morphology | **0.161 (0.038, 0.283)** | -2.278 (-3.992, -0.542) | 0.977 (0.717, 1.3) | 0.309 (0.026, 0.541) |
| Nitrogen 15 Isotope | **0.122 (0.008, 0.238)** | -1.645 (-3.193, -0.08) | 0.86 (0.587, 1.211) | 0.321 (0.013, 0.58) |
| Phylogenetic Diversity | **0.144 (0.03, 0.258)** | -1.896 (-3.469, -0.331) | 0.902 (0.652, 1.218) | 0.302 (0.019, 0.54) |
| Tail Length | **0.184 (0.062, 0.307)** | -2.624 (-4.331, -0.924) | 0.989 (0.727, 1.321) | 0.352 (0.052, 0.573) |

| **Density** | **Beta** | **Intercept** | **sigma** | **$R^2$** |
|---|---|---|---|---|
| Body Size | 0.005 (-0.095, 0.106) | -0.156 (-1.583, 1.269) | 0.819 (0.581, 1.132) | 0.076 (0.001, 0.245) |
| Carbon 13 Isotope | -0.021 (-0.117, 0.073) | 0.363 (-0.904, 1.673) | 0.712 (0.461, 1.042) | 0.125 (0.001, 0.36) |
| Cranial Shape | 0.028 (-0.064, 0.118) | -0.46 (-1.734, 0.818) | 0.721 (0.505, 1.011) | 0.104 (0.001, 0.311) |
| Cranial Size | -0.022 (-0.082, 0.037) | 0.357 (-0.478, 1.209) | 0.471 (0.311, 0.708) | 0.13 (0.001, 0.373) |
| Dentary Shape | 0.012 (-0.075, 0.099) | -0.248 (-1.481, 1.013) | 0.69 (0.478, 0.969) | 0.09 (0.001, 0.273) |
| Dentary Size | -0.021 (-0.087, 0.046) | 0.461 (-0.483, 1.393) | 0.535 (0.354, 0.787) | 0.115 (0.001, 0.339) |
| Ear Length | 0.062 (-0.035, 0.158) | -0.93 (-2.271, 0.441) | 0.782 (0.551, 1.085) | 0.156 (0.002, 0.399) |
| External Proportions | **0.191 (0.073, 0.307)** | -1.899 (-3.573, -0.252) | 0.937 (0.673, 1.271) | 0.417 (0.084, 0.624) |
| Head Shape | 0.042 (-0.056, 0.138) | -0.633 (-1.99, 0.73) | 0.766 (0.536, 1.07) | 0.118 (0.001, 0.344) |
| Head-Body Length | -0.02 (-0.127, 0.087) | 0.261 (-1.258, 1.774) | 0.867 (0.624, 1.188) | 0.082 (0.001, 0.257) |
| Hind Foot Length | 0.05 (-0.06, 0.162) | -0.74 (-2.304, 0.814) | 0.906 (0.655, 1.228) | 0.107 (0.001, 0.32) |
| Isospace | 0.04 (-0.066, 0.149) | -0.594 (-2.09, 0.854) | 0.795 (0.527, 1.158) | 0.141 (0.001, 0.399) |
| Locomotor Mode | **0.344 (0.226, 0.46)** | -4.084 (-5.715, -2.473) | 0.961 (0.698, 1.296) | 0.671 (0.442, 0.776) |
| Mass | 0.037 (-0.069, 0.141) | -0.498 (-1.953, 0.972) | 0.813 (0.575, 1.131) | 0.106 (0.001, 0.32) |
| Morphology | **0.112 (0.015, 0.21)** | -1.459 (-2.833, -0.081) | 0.801 (0.571, 1.093) | 0.277 (0.012, 0.522) |
| Nitrogen 15 Isotope | 0.058 (-0.038, 0.152) | -0.97 (-2.255, 0.321) | 0.712 (0.464, 1.05) | 0.208 (0.003, 0.487) |
| Tail Length | **0.07 (0.015, 0.125)** | -1.296 (-2.071, -0.5) | 0.43 (0.279, 0.648) | 0.395 (0.033, 0.628) |

**Supporting Table 4**: Differences in the Standardized Effect Sizes of Phylogenetic Diversity between communities when the distantly related *Haeromys minahassae* is included in the estimation. *H. minahassae* (Hm) is found in Ambang, Buliohuto, Dako, Gandang Dewata, Nokilalaki, and Torompupu. Removing Hm from the analysis reduces the PD SES in all communities where it is present (except species-rich Gandang Dewata) and increases the PD SES in all communities where it is absent (except species-poor Buliohuto). Latimojong becomes the community with the highest PD SES when Hm is removed, and Ambang the lowest. However, the effect of species richness on PD SES scores, estimated using Bayesian linear regression, remains nearly the same in both cases (result in main text).

| Community | No. Species with Hm | No. Species without Hm | PD SES with Hm | PD SES without Hm |
|---|---|---|---|---|
| Ambang | 13 | 12 | -0.18 | -1.59 |
| Bawakaraeng | 7 | 7 | -1.34 | -1.42 |
| Buliohuto | 10 | 9 | 0.57 | -0.30 |
| Dako | 12 | 11 | 1.09 | 0.85 |
| Gandangdewata | 23 | 22 | 1.26 | 1.45 |
| Katopasa | 12 | 12 | -2.05 | -1.36 |
| Latimojong | 13 | 13 | -0.24 | 1.77 |
| Nokilalaki | 16 | 15 | 0.49 | 0.08 |
| Torompupu | 15 | 14 | 0.98 | 0.78 |
